# Estimating the effective sample size in association studies of quantitative traits

**DOI:** 10.1101/2019.12.15.877217

**Authors:** Andrey Ziyatdinov, Jihye Kim, Dmitry Prokopenko, Florian Privé, Fabien Laporte, Po-Ru Loh, Peter Kraft, Hugues Aschard

**Affiliations:** Department of Epidemiology, Harvard T.H. Chan School of Public Health, Boston, MA, USA; Genetics and Aging Unit and McCance Center for Brain Health, Department of Neurology, Massachusetts General Hospital, Boston, MA, USA; Harvard Medical School, Boston, MA, USA; National Centre for Register-Based Research, Aarhus University, Aarhus, 8210, Denmark; Centre de Bioinformatique, Biostatistique et Biologie Intégrative (C3BI), Institut Pasteur, Paris, France; Division of Genetics, Department of Medicine, Brigham and Women’s Hospital and Harvard Medical School, Boston, MA, USA; Program in Medical and Population Genetics, Broad Institute of MIT and Harvard, Cambridge, MA, USA

## Abstract

The effective sample size (ESS) is a quantity estimated in genome-wide association studies (GWAS) with related individuals and/or linear mixed models used in analysis. ESS originally measured relative power in family-based GWAS and has recently become important for correcting GWAS summary statistics in post-GWAS analyses. However, existing ESS approaches have been overlooked and based on empirical estimation. This work presents an analytical form of ESS in mixed-model GWAS of quantitative traits, which is derived using the expectation of quadratic form and validated in extensive simulations. We illustrate the performance and relevance of our ESS estimator in common GWAS scenarios and analytically show that (i) family-based studies are consistently underpowered compared to studies of unrelated individuals of the same sample size; (ii) conditioning on polygenic genetic effect by linear mixed models boosts power; and (iii) power of detecting gene-environment interaction can be substantially gained or lost in family-based designs depending on exposure distribution. We further analyze UK Biobank dataset in two samples of 336,347 unrelated and 68,910 related individuals. Analysis in unrelated individuals reveals a high accuracy of our ESS estimator compared to the existing empirical approach; and analysis of related individuals suggests that the loss in effective sample size due to relatedness is at most 0.94x. Overall, we provide an analytical form of ESS for guiding GWAS designs and processing summary statistics in post-GWAS analyses.

## Introduction

Genome-wide association studies (GWAS) have identified thousands of genetic variant-trait associations, improving our understanding of genetic architecture of complex traits and diseases^1^. Most GWAS use linear regression performed in a sample of unrelated individuals, because statistical tests are computationally fast and have well-known analytical properties^2^. Importantly, post-processing methods based on GWAS summary statistics also assume a linear regression data model. These post-GWAS methods, including meta-analysis^3^, fine-mapping^4^, partitioning heritability^5,6^ and polygenic risk prediction^7^, are valuable resources to follow up GWAS findings and gain insights about the genetic architecture. However, one needs to estimate the effective sample size (ESS) to correct for potential sample relatedness and/or account for linear mixed models used to generate summary statistics^4,5^. Some methods recognized the problem of ESS estimation and incorporated data-driven approaches to handle ESS^4,5^. Ignoring the correction by ESS can produce misleading results such as overestimation of heritability enrichment^5^ and inaccurate fine-mapping of causal variants^4^.

Nowadays, modern cohorts consist of combined samples of unrelated and related individuals, for instance UK Biobank^8^, that poses a challenge to both GWAS and post-GWAS analyses. In the GWAS context, linear mixed moles (LMM) have been established as an effective alternative to linear regression (LR) performed on a subsample of unrelated individuals: LMM can be applied to the whole sample (related individuals retained)^9^, account for family or cryptic relatedness (control of spurious associations)^10^ and condition on the polygenic signal (power gain)^11^. Despite these well-known advantages of using LMM in GWAS^11^, works on optimizing computational algorithms and determining analytical properties remain an active area of research^9,11–13^. Here we are interested in an analytical expression for power of LMM association tests and their relative performance against LR. For ease of interpretation we use the ESS multiplier, defined as a ratio of the non centrality parameters (NCP) between the two tests and served as a measure of relative power^14^. We define a baseline scenario: testing the genetic effect on a quantitative trait by LR in a sample of unrelated individuals. The NCP for LR is known to be directly proportional to the sample size and the variance explained by genetic variant^2^. We next can derive the NCP for a variety of scenarios using LMM tests and analytically compare them to the baseline.

The scope of scenarios covered in this work is limited to three particular comparisons, which were previously discussed but, in our opinion, require an additional analytical revision in terms of relative power. These scenarios represent different study designs (unrelated/related individuals), association models (LR/LMM) and parameters of interest (marginal genetic/gene-environment interaction effects). First, we aim at providing an analytical solution to quantify the impact of having related rather than unrelated individuals in a sample^9,15^. Intuitively, having related individuals results in lowering the power, as related pairs harbor overlapping phenotypic and genetic information^15^. Related works provided an analytical solution of ESS only for special cases such as sibling pairs^16^. Second, we revisit the impact of using LMM in association study of unrelated individuals, where the polygenic signal is modeled as a random effect via the genetic relationship matrix. Previous works were focused on the distribution of test statistic^2,11^ and proposed to estimate ESS empirically using top statistic^5,9^. Third, we tackle association studies of gene-environment interactions^17^ and examine how family resemblance in related individuals affects the power of detecting interaction effects using LMM. Related works empirically evaluated different family-based designs to improve power^18,19^, but the complete analytical derivation for interaction test statistic is available only for LR applied to unrelated individuals^17^.

This work presents a formal framework to compare the relative performance of different LMM association tests in respect to the baseline LR test. The manuscript is organized as follows. We first derive approximations of NCP for LMM tests and use them to further derive the ESS multiplier. We then demonstrate the validity of our multiplier through extensive simulations and real data analysis in UK Biobank^8^. We point out factors that influence the relative power: family structure, variance explained by LMM components such as heritability, and distribution of environmental exposure when testing for gene-environment interactions.

## Methods

### Linear models

We consider a linear mixed model (LMM) and derive the Wald test statistic of association between a genetic variant and a quantitative trait. We further derive the linear regression (LR) statistic as a special case of LMM statistic.

Let denote *N* is the number of individuals, *M* is the number of genetic variants, *y* is a *N* ×1 vector of trait, *W* is a *N* ×*M* matrix of genetic variants and *w* is a *N* ×1 vector of the genetic variant tested, i.e. a column in *W*. We assume that the vector *y* and the columns in matrix *W* are standardized to have zero mean and unit variance, and there are no other covariates. We then model *y* by a multivariate normal distribution.

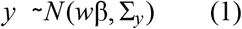

where β is the standardized effect size of variant *w*, and Σ_*y*_≡*cov*(*y*) is the *N* ×*N* covariance matrix of trait across *N* individuals.

If the covariance matrix Σ_*y*_ is known, β can be estimated using Generalized Least Squares (GLS)^20,21^. Then the Wald statistic is defined as 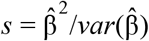, and it is compared to 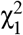 distribution under the null hypothesis of no association, β = 0. Thus, the LMM statistic is expressed as follows^12,20,21^.

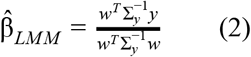

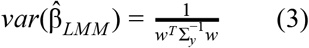

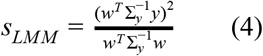

The LR statistic has a simpler form compared to Equations (2)–(4), considering that 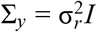 and *w* is standardized (*w*^*T*^*w* = *N*) and assuming 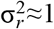, as the vector *y* is standardized and the variance captured by genetic variant is negligibly small.

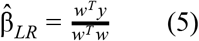

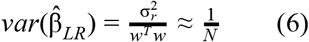

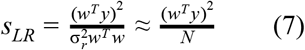

### Testing gene-environment interaction

To study the gene-environment interaction effect on quantitative trait *y*, the linear model in Equation (1) is expanded by including two *N* ×1 vectors: one vector *d* of environmental exposure, and another vector *v*≡*w* * *d* of gene-environment interaction obtained by element-wise multiplication of the two vectors *w* and *d*.

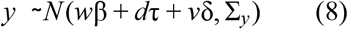

where β, **τ** and **δ** denote the effect sizes of genetic variant, exposure and interaction, respectively. We again assume that all the three vectors of covariates are standardized to have zero mean and unit variance, and there are no other covariates.

Under an assumption that two random variables of genotype and environmental exposure are generated independently, the *standardized* interaction effect δ can be evaluated independently from the two main effects β and τ ^17^. Thus, the test statistic for gene-environment interaction looks the same as in Equations (2)–(7) with replacement of *w* by *v*.

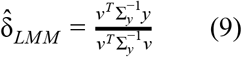

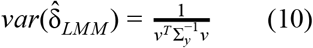

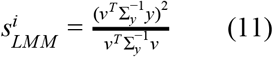

### Estimating trait covariance

The covariance structure of *y* is generally unknown, but Equations (1) and (8) can be extended to further specify covariance components. The expression for *y* can be written as follows.

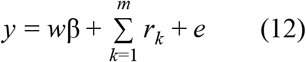

where *m* vectors of random effects, 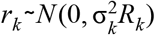, and residual errors, 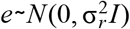, are assumed mutually uncorrelated and multivariate normally distributed. The covariance of each vector of random effects is parametrized with constant matrix *R*_*k*_ and scaled by the scalar parameter 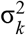, referred to as variance components. Marginalizing over vectors of random effects from Equation (12) gives a multivariate normal distribution of *y* with covariance given as follows.

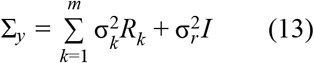

Both fixed effect β and variance components 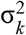 and 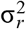, are model parameters. Variance components are typically estimated by restricted maximum likelihood (REML)^22^, because the REML approach produces unbiased estimates by adjustment for the loss in degrees of freedom due to the fixed effect covariates. To compute the association test statistic in Equations (4) and (7), we replace the true trait covariance by its estimate.

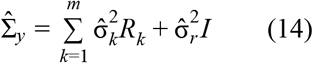

### Relative Power and Effective Sample Size

Under the alternative hypothesis, the non-centrality parameter (NCP) quantifies the statistical power for a given effect size β.

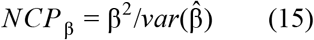

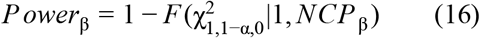

where α is the type I error rate, *F* (χ^2^|*df*, *NCP*) is the cumulative distribution function for the non-central χ^2^ distribution with *df* degrees of freedom and non-centrality parameter *NCP*. The quantity 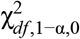 is the inverse of F or the quantile of the non-central χ^2^ distribution.

To introduce the concept of relative power and effective sample size (ESS), consider two association study designs based on unrelated individuals and related individuals in families. Both studies have the same sample size N, and one is interested to know which design is more powerful to detect a genetic variant with effect size β. For two association models, LR for unrelated individuals and LMM for related individuals, we derive the ratio of the two corresponding NCPs as defined in Equations (3) and (6)

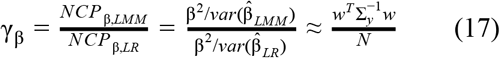

This ratio γ_β_, the ESS multiplier, is a measure of relative power and, by default, gives the ratio of the sample sizes needed for two study designs to yield the same variance of estimate^15^. Note that the ESS multiplier is similar to the asymptotic relative efficiency of two tests, say one likelihood to another, for measuring a parameter θ: it is given by the ratio of the inverse asymptotic estimates for the variance of 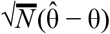^14^. In this work we aim to simplify the numerator part of ratio γ_β_ and propose approximations, as described in the next section.

Alternatively, an empirical estimator of the ESS multiplier, γ_*e*_, can be used when the analytical form of multiplier is unknown. An empirical solution has been proposed using test statistics at a subset of variants with the strongest association^5,9^. Consider two association studies in a sample of unrelated individuals, one being performed with LR and the other one with LMM. We derive the ratio of statistics computed by LMM and LR at *M*_*t*_ top associated variants, for example, at genome-wide significant variants based on LMM results. The empirical multiplier γ_*e*_ for these variants has the following form.

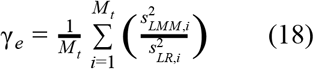

We note that the choice of top variants is subjective and, more importantly, the empirical multiplier takes average over ratios between association statistics rather than standard errors. The premise of this approach is based on the assumption that the estimated effect sizes at the top variants for LR and LMM are approximately the same and therefore cancel out in the ratio. When this assumption holds, the two estimators, γ_*e*_ and γ_β_, are equivalent.

### Approximations

Given the definition of the NCP in Equation (15), we compute the expected variance of the effect size estimate in Equation (3) by averaging 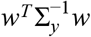 over genetic variants *w*, in order to obtain an analytical approximation for the NCP and power to detect a given effect size β. A similar computation is performed for NCP and power to detect gene-environment interaction effect size δ by averaging 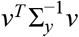 over interaction variables *v*. In particular, we approximate quadratic forms from LMM association models, 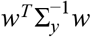 and 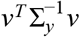, by their mean values, considering *w* and *v* as vectors of random variables and 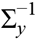 as a constant matrix of linear transformation.

First, we introduce the covariance matrix of genetic variant, Σ_*w*_≡*cov*(*w*), that convey genetic relatedness or pedigree structure of individuals. For unrelated individuals, Σ_*w*_ is the identity matrix. For related individuals in families, Σ_*w*_ is the expected kinship matrix, Σ_*w*_ = *K*, and is determined from pedigree information.

Second, we note that the covariance matrix of gene-environment interaction variable, Σ_*v*_≡*cov*(*v*), can be derived from Σ_*w*_ through the vector of environmental exposure, *d*, given in Equation (8). Briefly, we replace definition of *v* through element-wise multiplication of vectors *w* and *d* and introduce a matrix *E* = *diag*(*d*). Treating the matrix *E* as constant and *w* as a random vector, we obtain *cov*(*Ew*) = *E*Σ_*w*_*E*^*T*^. Next, we simplify the last expression by taking into account that the matrix *E* is diagonal. Defining a new matrix *D* and using the Hadamard product operator (°), we obtain the final form of Σ_*v*_.

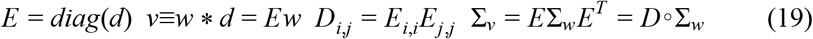

While the case of unrelated individuals with Σ_*w*_ = *I* is trivial and gives Σ_*w*_ = *diag*(*D*), we denote a special kinship matrix *K*_*D*_ for related individuals when Σ_*w*_ = *K*.

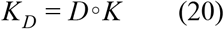

A numerical example of matrices *E*, *D*, *K* and *K*_*D*_ for nuclear families and binary exposure is provided in Supplementary Material.

Third, we approximate quadratic forms by their expected values. If *X* is a vector of random variables with mean μ and (nonsingular) covariance matrix Σ, then the quadratic form is a scalar random variable with mean expressed as follows.

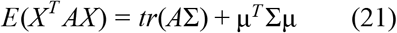

The variables *w* and *v* are standardized to have zero mean, then we obtain approximations.

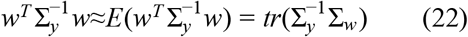

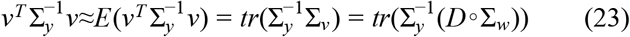

Fourth, we consider several LMM-based scenarios with particular structure of covariance matrices Σ_*y*_, Σ_*w*_ and Σ_*v*_ (see Tables 1 and 2). For each of these scenarios we propose further approximations of Equations (22) and (23) using known relationships between the trace operator and eigen-value decomposition outlined in Supplementary Material.

**Table 1:** Scenarios and covariance matrices for testing marginal genetic effect.

| Scenario | Model | Study design (Individuals) | $\Sigma_y$ | $\Sigma_w = K$ |
| --- | --- | --- | --- | --- |
| Unrelated | LR | Unrelated | $\sigma_r^2 I$ | $I$ |
| Families | LMM | Related | $\sigma_a^2 K + \sigma_r^2 I$ | $K$ |
| Unrelated+Grouping | LMM | Unrelated | $\sigma_f^2 F + \sigma_r^2 I$ | $I$ |
| Unrelated+GRM | LMM | Unrelated | $\sigma_g^2 G + \sigma_r^2 I$ | $I$ |

**Table 2:** Scenarios and covariance matrices for testing gene-environment interaction effect.

| Scenario | Model | Study design (Individuals) | $\Sigma_y$ | $\Sigma_v = K_D = D \circ K$ |
| --- | --- | --- | --- | --- |
| Unrelated | LR | Unrelated | $\sigma_r^2 I$ | $diag(D)$ |
| Families | LMM | Related | $\sigma_a^2 K + \sigma_{ai}^2 K_I + \sigma_r^2 I$ | $D \circ K$ |
| Unrelated+Grouping | LMM | Unrelated | $\sigma_f^2 F + \sigma_r^2 I$ | $diag(D)$ |
| Unrelated+GRM | LMM | Unrelated | $\sigma_g^2 G + \sigma_{gi}^2 G_I + \sigma_r^2 I$ | $diag(D)$ |

### Scenarios

We consider four GWAS scenarios to compare their relative power (Tables 1 and 2). Scenarios differ by study design, whether the data is collected for genetically unrelated or related individuals in families^15^. Additionally, studies of unrelated individuals vary by association models, LR or LMM. When analyzing unrelated individuals using LMM and testing for marginal genetic effect, we limit our comparisons to LMM with a single random effect, which is either a grouping factor, e.g. household, or a polygenic effect with genetic relationship matrix (GRM)^11^. In all scenarios, the vector of trait *y* is standardized, so that the sum of variance components in Σ_*y*_ (scalars 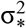) is equal to 1. The parameter 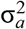 denotes the additive heritability in family-based study. The other similar parameter 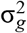 stands for the heritability explained by genetic variants in study of unrelated individuals.

### Data simulations

In simulations, we generate a quantitative trait from a multivariate normal distribution with variance components specified in Tables 1 and 2. In power analysis testing marginal genetic effect, we draw β so that the genetic variant explains ≈ 0.1% of trait variance. In power analysis testing gene-environment interaction effect, we draw δ so that the (standardized) gene-environment interaction term explains ≈ 0.1% of trait variance (standardized main genetic and environmental effects each explains 0.1% of trait variance). See Supplementary Material for more details.

In simulations of related individuals, we generate data for nuclear families with 2 parents and 3 offspring, if not specified otherwise. Accordingly, the kinship matrix *K* is added as a component of Σ_*y*_ for controlling the family structure in trait covariance. A special matrix *K*_*I*_ is also included in Σ_*y*_ when testing for gene-environment interaction^23^. Note that matrices *K*_*D*_ in Equation (19) and *K*_*I*_ in ref.^23^ are different, although both are derived from the kinship matrix *K* and realized exposure variable. In simulations of unrelated individuals with a grouping factor, each group consists of 5 individuals. Thus, the variance-covariance matrix, *F*, is a Kronecker product of block and diagonal matrices, where each block matrix is a 5×5 matrix of ones.

### Analysis of UK Biobank

In analysis of 336,347 UK Biobank unrelated individuals, we perform two LR- and LMM-based GWAS and then estimate the ESS multiplier between the two studies (rows 1 and 4 in Table 1). We follow a computationally efficient approach of low-rank LMM^24–26^, where LMM has a single random genetic effect with genetic relatedness matrix (GRM) constructed on a subset of top 1,000 SNPs, as described in another UK Biobank application^26^. These 1,000 SNPs are selected from top clumped LR associated SNPs using plink 2.0 (r^2^<0.1)^27^. The analysis is restricted to 336,347 British-ancestry unrelated individuals passing principal component analysis filters and having no third-degree or closer relationships^8^; 619,017 high-quality genotyped autosomal SNPs with missingness <10% and minor allele frequency (MAF) >0.1%^9^; six anthropometric traits, body mass index (BMI), height, hip circumference (HIP), waist circumference (Waist) and waist-to-hip ratio (WHR). To account for population structure, 40 principal components (PC) are included as covariates. Note that the performed low-rank LMM GWAS is not the most optimal strategy^11^, but it is sufficient to compare the relative performance of ESS multipliers.

In analysis of 68,910 UK Biobank related individuals, we select 40,231 related pairs with at least the second-degree relatedness to compute the ESS multiplier (rows 2 in Table 1). Kinship coefficients are empirically estimated from genotype data and allow to further split related pairs into categories (monozygotic twins, parent-offspring, full siblings and second-degree relatives), as described in ref. ^8^ and summarized in Supplementary Table S1.

### Efficient computation

Computation of quantities in Equations (22) and (23) requires inverting the trait covariance matrix 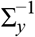. This is prohibitive in large datasets, so we developed several solutions to mitigate the computational burden. When Σ_*y*_ is dense in analysis of unrelated individuals, we follow the low-rank LMM approach implemented in a specially developed R package (github.com/variani/biglmmz). Our package is built on the R packages bigstatsr and bigsnpr with statistical methods for large genotype matrices stored on disk^28^. When Σ_*y*_ is sparse in analysis of related individuals, we apply special linear algebra methods for sparse matrices implemented in the R package Matrix; this approach was recently proposed for biobank-scale association studies^29^. In both analytical derivations and analysis of family-based data, we exploit the block structure of kinship matrices when it is possible.

## Results

### Analytical estimators for the effective sample size multipliers

We analytically derived γ_β_, the ESS multiplier of LMM against LR across the four scenarios described in Table 1. Recall a genetic variant *w* with effect β on a quantitative trait *y* with covariance matrices of trait and genetic variant Σ_*y*_ and Σ_*w*_, respectively. Using approximations given in Equation (22) (Methods), the relative power between LR and LMM tests can be approximated as follows:

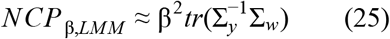

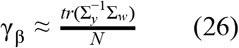

Expanding Equation (26) for each scenario in Table 1 and using components of Σ_*y*_ and the form of Σ_*w*_, we next obtain (see Supplementary Material):

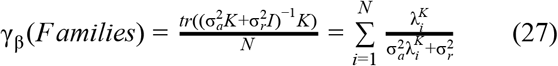

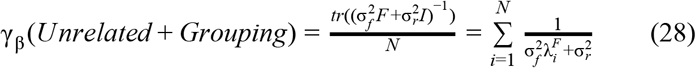

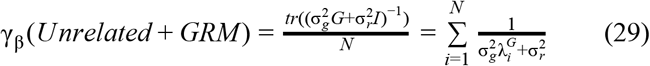

The multiplier for Families can be further simplified, for example, for related-pairs designs. If *s* is the number of related pairs within each family and r is the relatedness, then we obtain (see Supplementary Material):

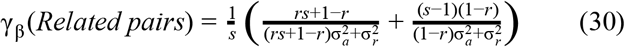

Finally, we similarly derived the NCP parameter for power to detect gene-environment interaction effect δ (Table 2). Given that the covariance matrices of trait and interaction variable are Σ_*y*_ and Σ_*v*_ = Σ_*w*_∘*D*, respectively, and the matrix *D* is defined in Equation (18), we obtain the approximation:

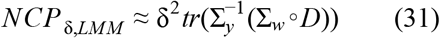

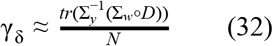

We also validated our approximations in Equations (25) and (31) through simulations (see Supplementary Figures S1-5).

### Testing marginal genetic effect

#### Power loss in related individuals

We examined the relative power for scenario Families (Table 1) by varying the heritability parameter 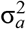. The multiplier γ_β_ for Families is strictly lower than 1 at all values of heritability except extreme values of 0 and 1 (blue lines on Figure 1a-b). The amount of power loss also depends on the structure of the matrices Σ_*y*_ and Σ_*w*_ = *K*. For example, the kinship matrix K for nuclear families with larger number of offspring leads to a greater loss, as K becomes more dense (Supplementary Figure S6). Similarly in studies of related pairs, monozygotic twin pairs show the power loss up to 50%, while decrease in power for pairs of siblings or cousins is moderate (Supplementary Figure S7).

**Figure 1:**
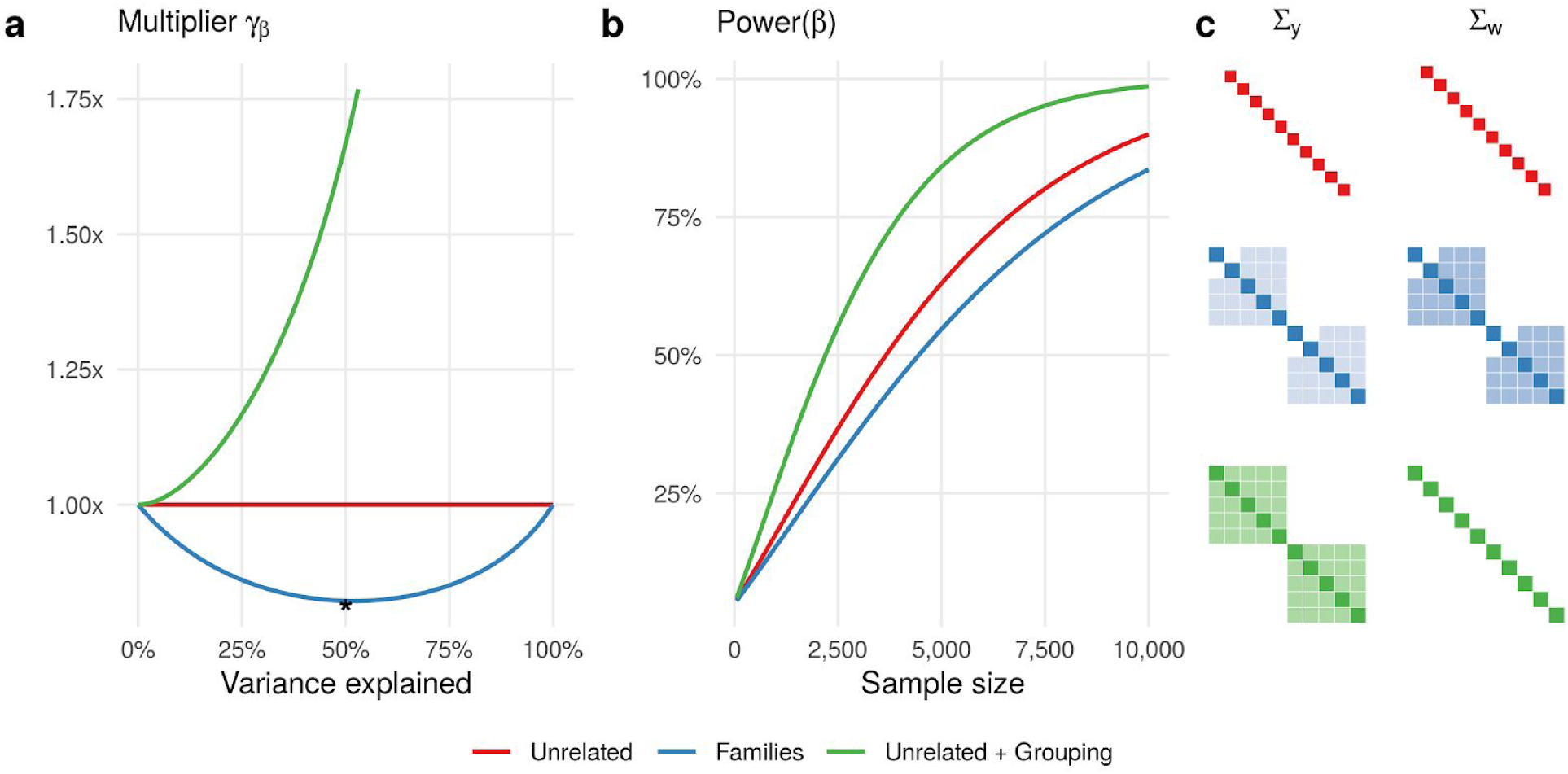
Relative power of detecting marginal genetic effect *β*. (a) The ESS multiplier γ_β_ is less than one for Families and greater than one for Unrelated+Grouping compared to the baseline scenario Unrelated. The amount of variance explained by random effect (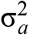 or 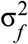) varies from 0% to 100%. (b) The power of detecting *β* increases with the sample size at different rates for Unrelated, Families and Unrelated+Grouping. The random effect and genetic variant explain 50% and 1% of trait variance, respectively. (c) The covariance matrices of trait and genetic variant *Σ*_y_ and *Σ*_w_ (used to compute γ_β_) are depicted when 50% of trait variance is explained by random effect.

The performance of multiplier for scenario Families is quantitatively described by formula in Equation (26), in which the trace operator is applied to product of two matrices 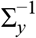 and Σ_*w*_ = *K*. To gain an intuition about the power loss for Families, we depict the covariance matrices Σ_*y*_ and Σ_*w*_ at 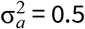 (Figure 1c). Off-diagonal non-zero entries of Σ_*w*_ (the double kinship coefficient, 0.5) are always lower than matched off-diagonal entries of 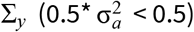, that explains why the multiplier is smaller than one.

#### Power gain by reducing residual variance

We varied the amount of variance explained by grouping factor 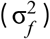 for scenario Unrelated+Grouping (Table 1) and observed the change in relative power. In contrast to Families, the gain in power for Unrelated+Grouping compared to Unrelated is consistent and increases as more variance is explained (green lines on Figure 1a-b). The observed increasing trend trivially follows from Equations (26) and (28) if one considers the trace operation 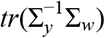 and takes into account that Σ_*w*_ = *I*. Thus, having individuals genetically unrelated (Σ_*w*_ = *I*) and explaining additional variance by a random effect is equivalent to a reduction in residual variance by including covariates, for example, using dummy variables from the grouping factor in scenario Unrelated+Grouping.

We further note that two scenarios Unrelated+Grouping and Unrelated+GRM (Table 1) are conceptually identical, because individuals are genetically unrelated. This implies that the observed trends on Figure 1 for Unrelated+Grouping are directly transferable to Unrelated+GRM. We confirmed this statement by simulations for Unrelated+GRM (Supplementary Material).

#### Modest power gain by low-rank LMM in UK Biobank unrelated individuals

Applying low-rank LMM to 336,348 UK Biobank unrelated individuals, we achieved a modest power gain with the maximum of 1.2x for height (Figure 2). Apart from boosting power, we revealed a high accuracy of our analytical multiplier γ_β_ compared to the empirical multiplier γ_*e*_. To get the true value of multiplier, we used the observed ratio of squared standard errors from LR and LMM tests (dark grey bars on Figure 2a but not on Figure 1b). We next compared the two multipliers γ_β_ and γ_*e*_ and observed that the multiplier γ_β_ (red bars) accurately approximates the observed ratio (Figure 2a). The empirical multiplier γ_*e*_ (beige bars on Figure 2b but not on Figure 2a), which is based on ratios of test statistics rather than standard errors, consistently underestimates the same observed ratio for all six traits (Figure 2b). The downward bias of γ_*e*_ is in agreement with our results on simulated data for Unrelated+GRM scenario (Supplementary Material).

**Figure 2:**
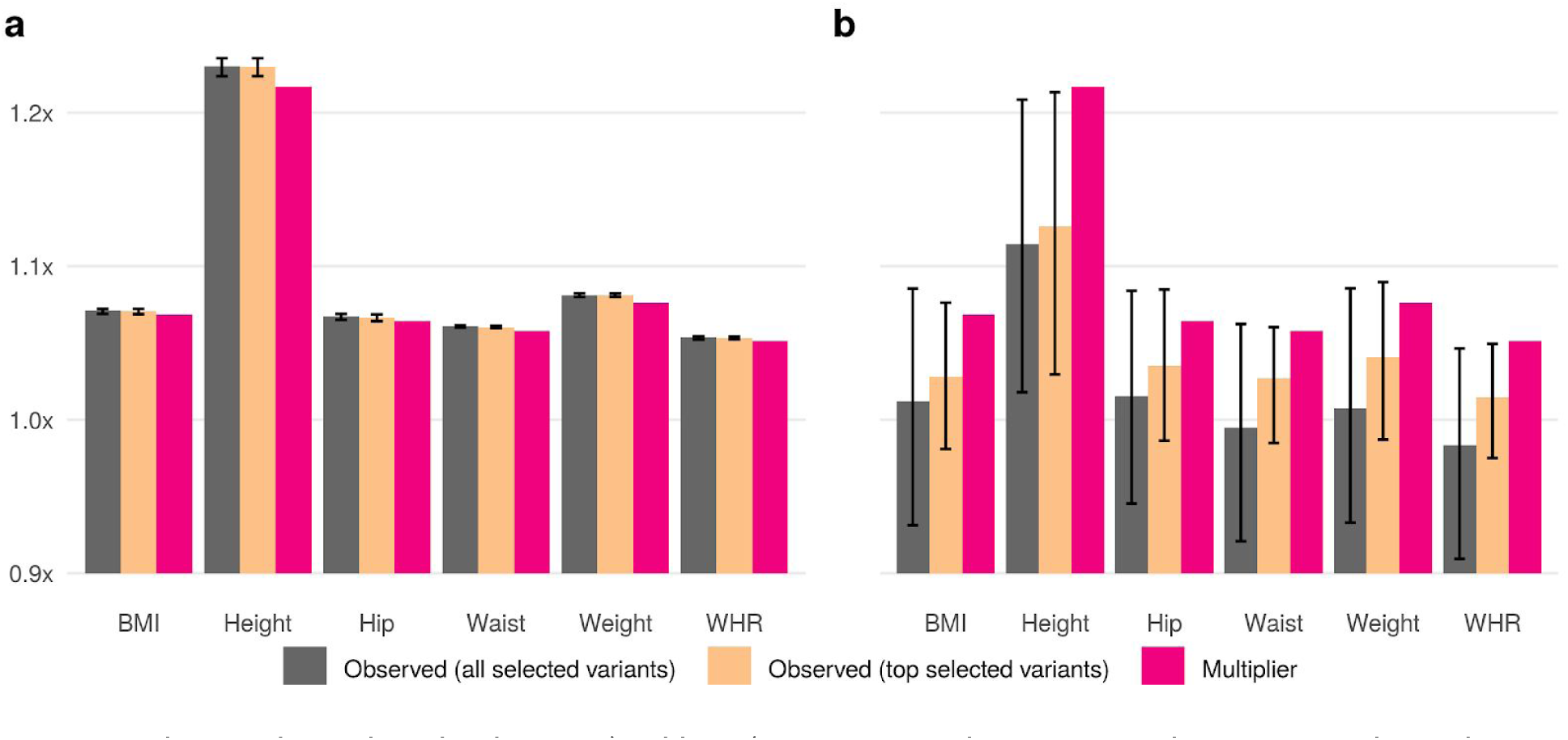
The analytical multiplier (red bars) is compared to empirical γ_β_ estimators based on (a) ratios of squared standard errors and (b) ratios of squared test statistic. Association studies of six anthropometric traits are performed using LR and low-rank LMM in 336,347 UK Biobank unrelated individuals. Empirical estimators are computed using either all 1,000 variants selected for low-rank LMM (dark grey bars) or a subset of 1,000 selected variants (significant in LMM, P_LMM_ < 1×10^−5^, and nominally significant in LR, P_LR_ < 0.05) (beige bars). Heights of dark grey and beige bars represent mean values, while error bars range from 1st to 3rd quartiles.

#### Small power loss in UK Biobank related individuals

We obtained estimates of the ESS multiplier γ_β_ for several groups of UK Biobank related pairs: monozygotic twins, parent-offspring, full siblings and second-degree relatives. All together for 68,910 close relatives of up to the second degree, the maximum drop in the effective sample size 0.94 was observed at heritability 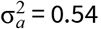. Considering the impact of relatedness in the whole UK Biobank sample, the multiplier 0.94 in related individuals is scaled to 0.99 in a combined sample of unrelated and related individuals. We also report minimum values of the multiplier stratified by groups of related pairs in Supplementary Table 2 and Supplementary Figure S8.

### Testing gene-environment interaction effect

#### Power gain or loss depends on realized environmental exposure and variance components

We explored the power gain for Families and Unrelated+Grouping scenarios over the baseline Unrelated when testing gene-environment interaction effect (Figure 3). The frequency of binary exposure was fixed to 0.6 for all three scenarios, but for Families we fixed the exposure status in such a way that two parents were unexposed and three offspring were exposed. Figure 3a-b shows that the ESS multiplier for Unrelated+Grouping and Families is always greater than 1 and increases as more variance is explained. This positive trend would remain for Unrelated+Grouping and Unrelated+GRM scenarios with other realizations of exposure, as the residual variance is simply reduced and individuals are unrelated. Contrary to Unrelated+Grouping and Unrelated+GRM, the power gain for Families was achieved through a particular realization of exposure and covariance matrices Σ_*y*_ and Σ_*v*_, as shown on Figure 3c.

**Figure 3:**
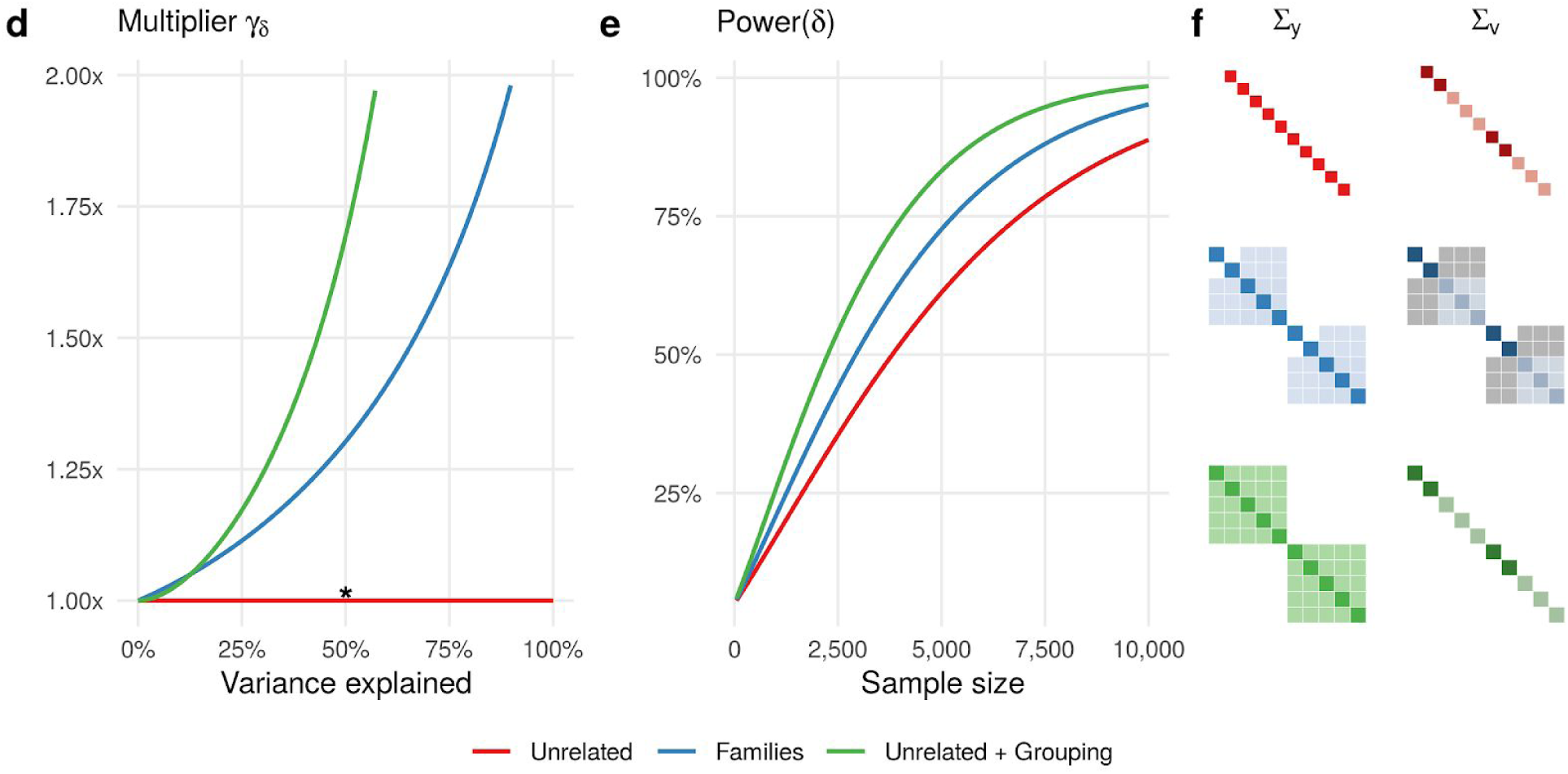
Relative power of detecting gene-environment interaction effect δ. The frequency of binary exposure is 0.6; the exposure status is fixed for Families, unexposed two parents and exposed three offspring. (a) The ESS multiplier γ_δ_ is greater than one for both Families and Unrelated+Grouping compared to the baseline scenario Unrelated. The amount of variance explained by random effects (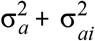 or 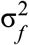) varies from 0% to 100%. (b) The power of detecting δ increases with the sample size at different rates for Unrelated, Families and Unrelated+Grouping. The random effects and genetic variant explain 50% and 1% of trait variance, respectively. (c) The covariance matrices of trait and interaction variable Σ_y_ and Σ_v_ (used to compute γ_δ_) are depicted when 50% of trait variance is explained by random effects. Colored gradient in entries of matrices denote quantitative differences for positive values, while grey-colored entries correspond to negative values. The ratio between 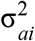 and 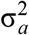 is fixed to 0.1; both genetic and environmental variables also explain 1% of trait variance in addition to 1% of interaction variable.

We next explored in more depth the relative power for Families as a function of exposure realization and interplay between covariance matrices Σ_*y*_ and Σ_*v*_ (Figure 4). In particular, we considered all possible realizations of binary exposure within families and also varied the composition of variance components in 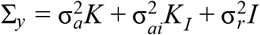 while fixing the total genetic variance, 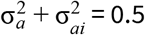. When the structure of Σ_*y*_ is fully defined by the kinship matrix K (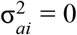, Figure 4, left panel), the multiplier is greater than 1.2 for all realizations of exposure and the most power gain 1.38 is achieved when all offspring are either exposed or unexposed. With increasing contribution of the environmental kinship matrix *K*_*I*_ into the structure of 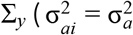 or 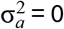, Figure 4, middle and right panels), the multiplier is getting closer to 1 and stands below 1 at 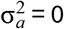. That occurs because the covariance matrices Σ_*y*_ and Σ_*v*_ become similar in their structure that leads to power loss. This phenomenon is similar to the analysis of Family scenario when testing marginal genetic effect (Figure 1, Supplementary Figures S6-8).

**Figure 4:**
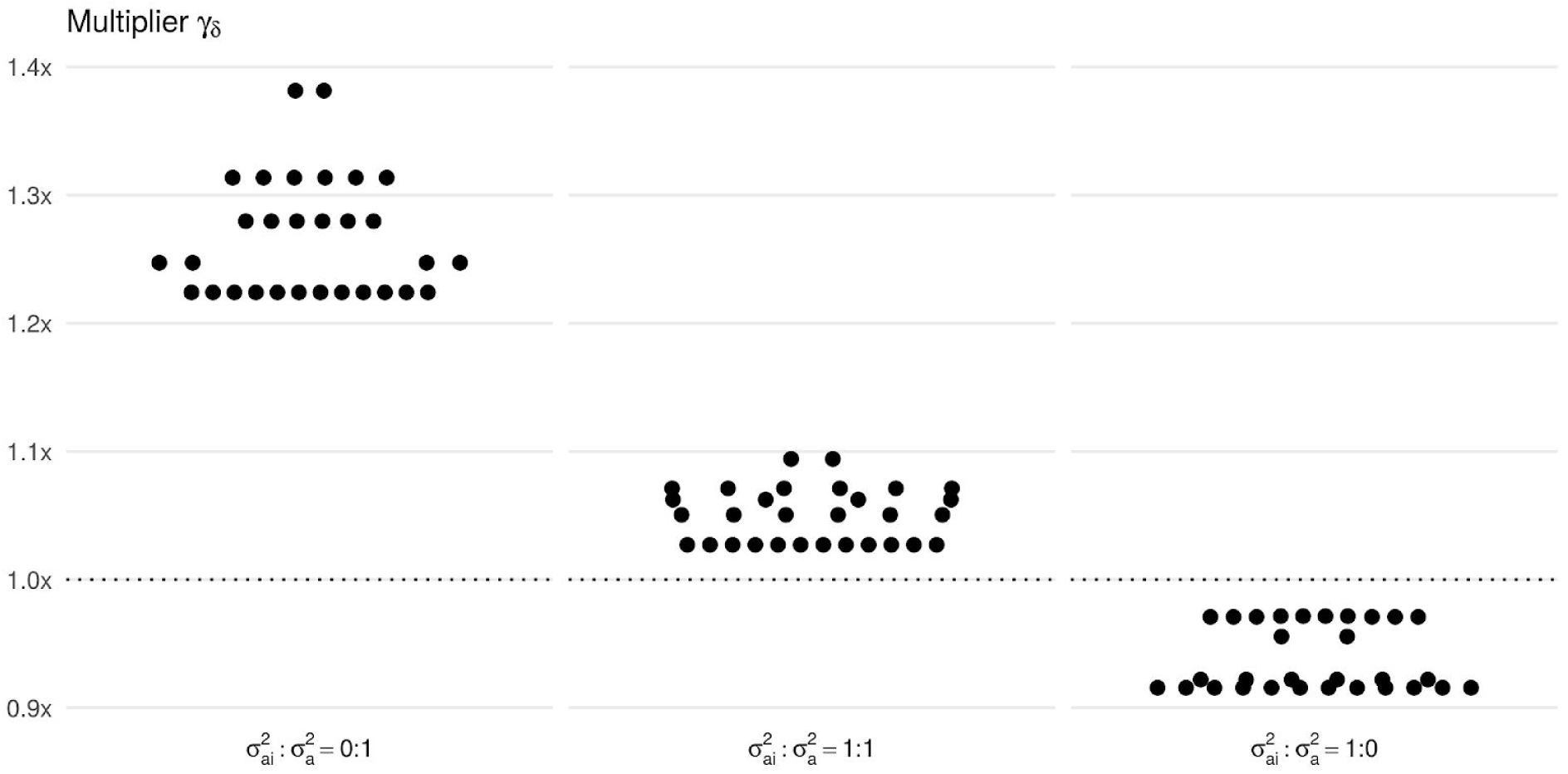
Relative power of detecting gene-environment interaction effect δ in nuclear families under different simulation settings. The ESS multiplier γ_δ_ is analytically computed (i) for all possible realizations of a binary exposure within a nuclear family with 2 parents and 3 offspring (dots in each panel) and (ii) for different ratios between 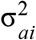 and 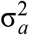 (three panels). The amount of trait variance jointly explained by random effects 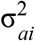 and 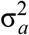 is fixed to 50%. The largest two values of multiplier on left and middle panels correspond to exposure realizations of exposed offspring/unexposed parents and exposed parents/unexposed offspring.

## Conclusions

Linear mixed models are increasingly used in genome-wide association studies. While of great benefit, the inference of mixed model parameters are more computationally and analytically complex than for standard linear regression models. To address analytical complexities specific to power, we derived the formula for NCP of mixed-model association tests, which is similar to NCP of linear regression (proportional to the sample size and variance captured by genetic variant), but it also incorporates trait covariance and genetic relatedness matrices. We further introduced the ESS multiplier, defined as a ratio between NCPs of two tests, and showed its performance in quantifying the relative power across common GWAS scenarios.

Compared to related works ^9,11^, we shifted the focus of our analysis from distribution of test statistics to distribution of standard errors of estimated effect sizes. While modeling distribution of statistics allows to distinguish confounding from polygenicity ^30^ and informs partitioning of heritability ^6^, errors terms are directly linked to the effective sample size and power. We covered common GWAS scenarios in our unified analytical framework, considering family-based studies as well as studies of unrelated individuals under association models with genetic or non-genetic random effects. Additionally, our analytical derivations were naturally extended to studies of gene-environment interactions.

Improving power of detecting gene-environment interaction by optimization of family-based designs is an attractive research area ^17–19^. We confirmed our hypothesis that study designs can be leveraged to increase power due to particular interplay between relatedness structure and realized environmental exposure. We showed a particular case of power gain in simulated nuclear families with exposed offspring and unexposed parents. These results suggest that exposures collected in cohorts with related individuals can be assessed in terms of gain or loss in power before conducting actual GWAS screening of gene-environment interactions.

There are still a number of methodological issues arising in GWAS that are also relevant to our work. Incomplete population stratification by PCs is documented for height in UK Biobank data analysis ^29,31^, and our multiplier can be affected by this phenomenon through the covariance matrices used to calculate ESS. In our UK Biobank analysis we noticed small discrepancies between estimates of the multiplier and observed ratios of squared standard errors. Furthermore, if variance components are misspecified, the distribution of test statistics can be inflated making power analysis invalid, especially in studies of related individuals ^10^. Also, we limited our analytical derivations to quantitative traits, and future work is warranted to extend our results to binary traits under the liability threshold model ^32–34^.

Overall, the proposed multiplier informs GWAS study designs in terms of power. Post-GWAS analyses need to consider reporting the effective sample size in summary statistics using our analytical form of multiplier.

## Acknowledgements

This work was supported by NIH grants R21HG007687 (NHGRI) and R01CA194393 (NCI). The research was conducted using the UK Biobank Resource under Application #16549.

