## Supplementary Material for "Estimating the effective sample size in association studies of quantitative traits"

2020-01-04

### Supplementary Material

#### Propositions

##### Quadratic form

If $\mathcal{X}$ is a vector of random variables with mean $\mu$ and (nonsingular) covariance matrix $\Sigma$, then the quadratic form $\mathcal{X}^T A \mathcal{X}$ is a scalar random variable:

$$E(\mathcal{X}^T A \mathcal{X}) = tr(A\Sigma) + \mu^T \Sigma \mu \tag{33}$$

$$Var(\mathcal{X}^T A \mathcal{X}) = 2tr(A\Sigma A\Sigma) + 4\mu A \Sigma A \mu \tag{34}$$

See ref.[1] for more details.

##### Linear transform of random vector

If $B$ is a constant matrix and $\mathcal{X}$ is a vector of random variables with mean $\mu$ and covariance matrix $\Sigma$, then $B\mathcal{X}$ is a vector of random variables:

$$E(B\mathcal{X}) = BE(\mathcal{X}) \tag{35}$$

$$Var(B\mathcal{X}) = BVar(\mathcal{X})B^T \tag{36}$$

The proof makes use of definitions of mean and variance.

##### Eigen-value decomposition (EVD)

If $K$ is the covariance matrix of size $n \times n$, that means $K$ is symmetric and positive semi-definite. Furthermore, EVD of $K$ is

$$K = QDQ^T = QDQ^{-1} \tag{37}$$

where $Q$ is an $n \times n$ orthogonal matrix of eigen-vectors and $D$ is a $n \times n$ diagonal matrix of eigen-values ($\lambda_K^i$ with $i$ from 1 to $n$).

EVD for the matrix inverse to $K$ is

$$K^{-1} = QD^{-1}Q^T \tag{38}$$

EVD for the matrix such as $V = aK + bI$, where $a$ and $b$ are scalars, $I$ is the $n \times n$ identity matrix, is

$$V = aK + bI = aQDQ^T + bI = aQDQ^T + bQIQ^T = Q(aK + bI)Q^T \tag{39}$$

##### Trace operator and eigen-value decomposition (EVD)

For the covariance matrix $K$ and the matrix $V = aK + bI$, we have the following series of equations in relation to the trace operator.

$$\begin{aligned} tr(K) &= \sum_{i=1}^{n} \lambda_K^i \\ tr(K^{-1}) &= \sum_{i=1}^{n} (\lambda_K^i)^{-1} \\ tr(V) = tr(aK + bI) &= \sum_{i=1}^{n} (a\lambda_K^i + b) \\ tr(V^{-1}) = tr((aK + bI)^{-1}) &= \sum_{i=1}^{n} (a\lambda_K^i + b)^{-1} \\ tr(V^{-1}K) = tr((aK + bI)^{-1}K) = tr((aI + bK^{-1})^{-1}) &= \sum_{i=1}^{n} (a + b(\lambda_K^i)^{-1})^{-1} \end{aligned} \tag{40}$$

In the last equation we used the following equality.

$$\begin{aligned} V^{-1}K &= (aK + bI)^{-1}K = (aK + bI)^{-1}(K^{-1})^{-1} \\ &= K^{-1}(aK + bI)^{-1} = (aI + bK^{-1})^{-1} \end{aligned} \tag{41}$$

#### Analytical derivation of multiplier $\gamma_\beta$ for families

The effective sample size (ESS) multiplier for family-based association studies is given in Equation (32) of the main text. We write down this result again.

$$\gamma_\beta = \frac{tr\left((K\sigma_a^2 + I\sigma_r^2)^{-1}K\right)}{N} \tag{42}$$

We can further rewrite the numerator using the relationship between the trace operator and eigen-value decomposition (EVD) of matrix $(K\sigma_a^2 + I\sigma_r^2)^{-1}K$ given in Equation (40) of this supplementary material.

$$\gamma_\beta = \frac{1}{N} \sum_{i=1}^{N} \frac{\lambda_i}{\lambda_i \sigma_a^2 + \sigma_r^2} \tag{43}$$

The assumption of $y$ being standardized leads to $\sigma_r^2 = 1 - \sigma_a^2$:

$$f(\sigma_a^2, \lambda) = \gamma_\beta = \frac{1}{N} \sum_{i=1}^{N} \frac{\lambda_i}{(\lambda_i - 1)\sigma_a^2 + 1} \tag{44}$$

##### Splitting $K$ into blocks of submatrics $K_s$

The sample generally consists of $N_s$ families such that there is no between-family genetic relatedness. Then the kinship matrix $K$ can be represented as a block matrix.

$$K = \begin{pmatrix} K_{s_1} & 0 & \dots & 0 \\ 0 & K_{s_2} & \dots & 0 \\ \dots & \dots & \dots & \dots \\ 0 & 0 & \dots & K_{s_{N_s}} \end{pmatrix}$$

Hence, the multiplier can be evaluated separately for each family using the kinship matrices $K_{s_k}$ for each family, where $k$ is from 1 to $N_s$, $s_k$ is the dimension of square matrix $K_{s_k}$.

For family-based designs such as related pairs (siblings or twins) the blocks are the same and the block matrix $K$ has the form.

$$K = \begin{pmatrix} K_s & 0 & \dots & 0 \\ 0 & K_s & \dots & 0 \\ \dots & \dots & \dots & \dots \\ 0 & 0 & \dots & K_s \end{pmatrix} \tag{45}$$

The matrix $K_s$ for sibling pairs is:

$$K_s = \begin{pmatrix} 1 & 0.5 \\ 0.5 & 1 \end{pmatrix}$$

The matrix $K_s$ for monozygotic twins is:

$$K_s = \begin{pmatrix} 1 & 1 \\ 1 & 1 \end{pmatrix}$$

In a general case we consider $s$ related pairs with the relatedness coefficients $r$, where $s$ is a positive integer and $r$ is from 0 (unrelated) to 1 (monozygotic twins).

$$K_s = \begin{pmatrix} 1 & r & \dots & r \\ r & 1 & \dots & r \\ \dots & \dots & \dots & \dots \\ r & r & \dots & 1 \end{pmatrix} \tag{46}$$

#### Eigenvalues of $K_s$

Denote that $\lambda_i$ are $s$ eigenvalues of $K_s$ matrix in Equation (46). These eigenvalues can be analytically calculated by representing the matrix $K_s$ as a weighted sum of two matrices (one of which is a diagonal matrix).

$$K_s = \begin{pmatrix} 1 & r & \dots & r \\ r & 1 & \dots & r \\ \dots & \dots & \dots & \dots \\ r & r & \dots & 1 \end{pmatrix} = \begin{pmatrix} r & r & \dots & r \\ r & r & \dots & r \\ \dots & \dots & \dots & \dots \\ r & r & \dots & r \end{pmatrix} + \begin{pmatrix} 1-r & 0 & \dots & 0 \\ 0 & 1-r & \dots & 0 \\ \dots & \dots & \dots & \dots \\ 0 & 0 & \dots & 1-r \end{pmatrix}$$

If $\lambda_1$ is the first eigenvalue and $\lambda_{-1}$ are the remaining $(s-1)$ eigenvalues, then we use results in Equation (39) and obtain.

$$\begin{aligned} \lambda_1 &= rs + (1-r) \\ \lambda_{-1} &= 0 + (1-r) = 1-r \end{aligned} \tag{47}$$

Therefore, the sum of eigenvalues of $K_s$ is simplified.

$$\sum_{i=1}^{s} \lambda_i = \lambda_1 + (s-1)\lambda_{-1} = [rs + (1-r)] + [(s-1)(1-r)]$$

| Related pairs | No. pairs, s | Relatedness, r | First eigenvalue, $\lambda_1$ | Other eigenvalues, $\lambda_{-1}$ |
|---|---|---|---|---|
| Monozygotic twins | s | 1 | s | 0 |
| Siblings | s | 1/2 | (s+1)/2 | 1/2 |
| Cousins | s | 1/4 | (s+3)/4 | 3/4 |

**Table**: Eigen values of submatrix of $K$ ($K_s$) with respect to the relatedness distribution.

#### Computing $\gamma_\beta$ for related pairs through EVD of $K_s$

To simplify the analytical form of multiplier in Equation (44), we reformulate it using the block-wise represenation of $K$ given in Equations (45) and (46).

$$f(\sigma_a^2, \lambda) = \frac{1}{s} \sum_{i=1}^{s} \frac{\lambda_i}{(\lambda_i - 1)\sigma_a^2 + 1} \tag{48}$$

Finally, we get the result in Equation (30) of the main text by summing weighted eigenvalues of $K_s$ derived in (47). Here we write the Equation (30) again.

$$\begin{aligned}
\gamma_\beta(\text{Related pairs}) &= \frac{1}{s}\left(\frac{rs+1-r}{(rs+1-r)\sigma_a^2+\sigma_r^2} + \frac{(s-1)(1-r)}{(1-r)\sigma_a^2+\sigma_r^2}\right) \\
&= \frac{1}{s}\left(\frac{rs+1-r}{(rs-r)\sigma_a^2+1} - \frac{(s-1)(1-r)}{(r\sigma_a^2+1}\right) \\
&= \frac{1}{s}\left(\frac{(s-1)r+1}{(s-1)r\sigma_a^2+1} - \frac{(s-1)(1-r)}{r\sigma_a^2+1}\right)
\end{aligned}$$

#### Minima of function $\gamma_\beta(\sigma_a^2)$

To minimize the function $f$ from Equation (48) with respect to $\sigma_a^2$ and get its extrema points, we only have to find the solution:

$$\begin{aligned}
\frac{\partial f(x,\lambda)}{dx} &= 0 \\
-\frac{1}{s}\sum_{i=1}^{s} \frac{\lambda_i^2 - \lambda_i}{((\lambda_i - 1)x + 1)^2} &= 0
\end{aligned}$$

Note that $f(0,\lambda) = f(1,\lambda) = 1$ except for twins case. For twins case, we have $f(0,(s,0,\ldots,0)) = 1$ and $\lim_{x\to 1} f(x,(s,0,\ldots,0)) = \frac{1}{s}$.

**Case 1**: Siblings (s, r = 1/2)

The solution is:

$$\begin{aligned}
&\frac{(\frac{s+1}{2})^2 - \frac{s+1}{2}}{((\frac{s+1}{2}-1)x+1)^2} = (s-1)\frac{1}{4(1-\frac{1}{2}x)^2} \\
&\frac{s^2-1}{((s-1)x+2)^2} = \frac{s-1}{(2-x)^2} \\
&(s-3)sx^2 + 8sx - 4s = 0 \\
&x = \frac{2(\sqrt{s+1}-2)}{s-3} \text{ if } s \neq 3 \\
&x = \frac{1}{2} \text{ if } s = 3
\end{aligned} \tag{49}$$

**Case 2**: Cousins (s, r = 1/4)

The solution is:

$$x = \frac{4(s-1)(\sqrt{3(s+3)}-4)}{3s^2-10s+7} \tag{50}$$

**Case 3**: Twins (s, r = 1)

Because only one eigen value is not null, the derivative of function $f$ with respect to $\sigma_a^2$ is:

$$\frac{\partial f(\sigma_a^2, \lambda)}{d\sigma_a^2} = -\frac{1}{s}\frac{s^2-s}{((s-1)\sigma_a^2+1)^2} < 0 \tag{51}$$

The monotonic decrease of the function $\gamma_\beta(\sigma_a^2)$ for twins is observed on Supplementary Figures S6 and S7.

### Relationship matrices $K$, $K_D$ and $K_I$

To study the gene-environment interaction effect $\delta$ on a quantitative trait $y$, Equation (8) in the main text outlines the model, $y \sim \mathcal{N}(w\beta + d\tau + v\delta, \Sigma_y)$, where $y$, $w$, $d$ are observed $N \times 1$ vectors of standardized trait, genetic variant and exposure, respectively; $v$ is a vector of interaction between genetic variant and exposure obtained by element-wise multiplication of $w$ and $d$; $\beta$, $\tau$, $\delta$ are the effect sizes; $\Sigma_y$ is the $N \times N$ covariance matrix of trait across $N$ individuals. The matrix $\Sigma_v \equiv K_D$ is a covariance matrix of the interaction variable $v$.

Table 2 in the main text, according the ref.[2], suggests to include two types kinship matrices $K$ and $K_I$ into $\Sigma_y$: $\Sigma_y = \sigma_a^2 K + \sigma_{ai}^2 K_I + \sigma_r^2 I$. Inclusion of the matrix $K_I$ controls for family structure in testing gene-environment interaction and protects from spurious associations (false positives).

Overall, the relationship matrices $K$, $K_D$ and $K_I$ define the behavior of test statistic in association studies of gene-environment interaction. So we would like to show how these matrices look like for a particular example of nuclear family.

#### Families

Consider a single nuclear family of 5 individuals, 2 parents and 3 offspring. The kinship matrix $K$ is:

$$K = \begin{pmatrix} 1 & 0 & 0.5 & 0.5 & 0.5 \\ 0 & 1 & 0.5 & 0.5 & 0.5 \\ 0.5 & 0.5 & 1 & 0.5 & 0.5 \\ 0.5 & 0.5 & 0.5 & 1 & 0.5 \\ 0.5 & 0.5 & 0.5 & 0.5 & 1 \end{pmatrix}$$

Consider next a binary environmental exposure $d$, which is drawn such that the first two individuals (parents) are unexposed and the last three individuals (offspring) are exposed to the environment. Thus, the frequency of binary exposure is $f = 0.6$.

$$d = \begin{pmatrix} 0 & 0 & 1 & 1 & 1 \end{pmatrix}$$

The matrix $K_I$ is computed by element-wise multiplication of $K$ and a special masking matrix $M$, which defines whether a pair of individuals have the same exposure status ($d$).

$$M = \begin{pmatrix} 1 & 1 & 0 & 0 & 0 \\ 1 & 1 & 0 & 0 & 0 \\ 0 & 0 & 1 & 1 & 1 \\ 0 & 0 & 1 & 1 & 1 \\ 0 & 0 & 1 & 1 & 1 \end{pmatrix}$$

$$K_I = M \circ K = \begin{pmatrix} 1 & 0 & 0 & 0 & 0 \\ 0 & 1 & 0 & 0 & 0 \\ 0 & 0 & 1 & 0.5 & 0.5 \\ 0 & 0 & 0.5 & 1 & 0.5 \\ 0 & 0 & 0.5 & 0.5 & 1 \end{pmatrix}$$

Equations (19) and (20) in the Methods section of main text introduce matrices $E$, $D$ and $K_D$. These matrices are further used to derive the test statistic in association studies of gene-environment interactions. See Equations (31) and (32) for the result of derivation.

We next show how the matrices $E$, $D$ and $K_D$ look like for our example of nuclear family and binary exposure. One can see further that the matrices are scaled by factor $f(1-f)$, because genetic and environmental exposure variables are standardized according to our association model, Equation (8).

The matrix $E$ is simply defined as $E = diag(d)$.

$$E = \frac{1}{\sqrt{f(1-f)}} \begin{pmatrix} -f & 0 & 0 & 0 & 0 \\ 0 & -f & 0 & 0 & 0 \\ 0 & 0 & (1-f) & 0 & 0 \\ 0 & 0 & 0 & (1-f) & 0 \\ 0 & 0 & 0 & 0 & (1-f) \end{pmatrix}$$

The matrix $D$ is defined by a property $D_{i,j} = E_{i,i}E_{j,j}$ for $i, j$ from 1 to $N$.

$$D = \frac{1}{f(1-f)} \begin{pmatrix} f^2 & f^2 & -f(1-f) & -f(1-f) & -f(1-f) \\ f^2 & f^2 & -f(1-f) & -f(1-f) & -f(1-f) \\ -f(1-f) & -f(1-f) & (1-f)^2 & (1-f)^2 & (1-f)^2 \\ -f(1-f) & -f(1-f) & (1-f)^2 & (1-f)^2 & (1-f)^2 \\ -f(1-f) & -f(1-f) & (1-f)^2 & (1-f)^2 & (1-f)^2 \end{pmatrix}$$

Further simplifying the notation, we obtain.

$$D = \begin{pmatrix} f/(1-f) & f/(1-f) & -1 & -1 & -1 \\ f/(1-f) & f/(1-f) & -1 & -1 & -1 \\ -1 & -1 & (1-f)/f & (1-f)/f & (1-f)/f \\ -1 & -1 & (1-f)/f & (1-f)/f & (1-f)/f \\ -1 & -1 & (1-f)/f & (1-f)/f & (1-f)/f \end{pmatrix}$$

The matrix $K_D$, which is the covariance matrix $\Sigma_v$ of the interaction variable $v$ in Equation (8), has the form.

$$\Sigma_v = K_D = D \circ K = \begin{pmatrix} f/(1-f) & 0 & -0.5 & -0.5 & -0.5 \\ 0 & f/(1-f) & -0.5 & -0.5 & -0.5 \\ -0.5 & -0.5 & (1-f)/f & 0.5(1-f)/f & 0.5(1-f)/f \\ -0.5 & -0.5 & 0.5(1-f)/f & (1-f)/f & 0.5(1-f)/f \\ -0.5 & -0.5 & 0.5(1-f)/f & 0.5(1-f)/f & (1-f)/f \end{pmatrix}$$

For illustration purposes, we replace $f$ by its value 0.6.

$$E = \frac{1}{\sqrt{0.24}} \begin{pmatrix} -0.6 & 0 & 0 & 0 & 0 \\ 0 & -0.6 & 0 & 0 & 0 \\ 0 & 0 & 0.4 & 0 & 0 \\ 0 & 0 & 0 & 0.4 & 0 \\ 0 & 0 & 0 & 0 & 0.4 \end{pmatrix}$$

$$D = \frac{1}{0.24} \begin{pmatrix} 0.36 & 0.36 & -0.24 & -0.24 & -0.24 \\ 0.36 & 0.36 & -0.24 & -0.24 & -0.24 \\ -0.24 & -0.24 & 0.16 & 0.16 & 0.16 \\ -0.24 & -0.24 & 0.16 & 0.16 & 0.16 \\ -0.24 & -0.24 & 0.16 & 0.16 & 0.16 \end{pmatrix}$$

$$\Sigma_v = K_D = \frac{1}{0.24} \begin{pmatrix} 0.36 & 0 & -0.12 & -0.12 & -0.12 \\ 0 & 0.36 & -0.12 & -0.12 & -0.12 \\ -0.12 & -0.12 & 0.16 & 0.08 & 0.08 \\ -0.12 & -0.12 & 0.08 & 0.16 & 0.08 \\ -0.12 & -0.12 & 0.08 & 0.08 & 0.16 \end{pmatrix}$$

Some elements of $K_D$ are negative, because genetic and environmental exposure variables are standardized in Equation (8). Figure 3c in the main text depicts these negative elements by gray color.

#### Unrelated individuals

We can check whether our family-based derivations of relationship matrices $K$, $K_D$ and $K_I$ are consistent with the case of unrealted individuals, for which the kinship matrix is the indentity matrix, $K = I$.

The vector $d$ and matrix $D$ are the same for unrelated individuals, but the covariance matrix has a simpler form, $\Sigma_v = diag(D)$.

$$d = \begin{pmatrix} 0 & 0 & 1 & 1 & 1 \end{pmatrix}$$

$$D = \begin{pmatrix} f/(1-f) & f/(1-f) & -1 & -1 & -1 \\ f/(1-f) & f/(1-f) & -1 & -1 & -1 \\ -1 & -1 & (1-f)/f & (1-f)/f & (1-f)/f \\ -1 & -1 & (1-f)/f & (1-f)/f & (1-f)/f \\ -1 & -1 & (1-f)/f & (1-f)/f & (1-f)/f \end{pmatrix}$$

$$\Sigma_v = D \circ I = diag(D) = \begin{pmatrix} f/(1-f) & 0 & 0 & 0 & 0 \\ 0 & f/(1-f) & 0 & 0 & 0 \\ 0 & 0 & (1-f)/f & 0 & 0 \\ 0 & 0 & 0 & (1-f)/f & 0 \\ 0 & 0 & 0 & 0 & (1-f)/f \end{pmatrix}$$

Further, we expect the multiplier $\gamma_\delta$ from Equation (32) to be one for unrelated individuals. Since we have $\Sigma_y = \sigma_r^2 I = I$; $\sigma_r^2 = 1$, we need to show that $tr(\Sigma_v) = N$ for unrelated individuals.

$$\gamma_\delta \approx \frac{tr(\Sigma_y^{-1}(\Sigma_v))}{N} = \frac{tr(diag(D)}{N} = \frac{(1-f)Nf/(1-f) + fN(1-f)/f}{N} = 1$$

### Previous works on estimating the relative power

Here we list previous works on relative power in association studies.

For association studies of related individuals in families, it was shown that the power is always lower for pairs of relatives when compared to unrelated individuals[3]. Consider a toy example of a single pair of unrelared individuals compared to a pair of related individuals ($N = 2$): a variant with true standardized effect size $\beta$ is tested. Given that $\rho$ is the phenotypic correlation and $r$ is the coefficient of relationship (e.g 0.5 for a pair of siblings), NCP for two related individuals can be derived using regression theory[3] and then compared to that of two unrelated individuals.

$$\begin{aligned} NCP_{sib.pair} &\approx 2\beta^2(1-\rho r)/(1-\rho^2) \\ NCP_{unrel.pair} &= 2\beta^2/(1-\beta^2) \approx 2\beta^2 \\ \gamma_\beta &= NCP_{sib.pair}/NCP_{unrel.pair} \approx 1-\rho(r-\rho) \end{aligned} \tag{52}$$

When all family resemblance is due to additive genetic effects ($\rho = h^2 r$, where $h^2$ denotes the heritability), the ratio in Equation (52) is always less than one. Further work derived the NCP parameter for a more general case of sibling pairs[4].

In studies that contain both unrelated and related individuals, power loss is due to discarding related pairs and limiting the analysis to a subsample of unrelated individuals. The power loss due to the sample size reduction was previously quantified analytically for twin studies[5] and empirically for biobank-scale association studies[6].

In family-based association studies of gene-environment interactions, different strategies of selecting family members for analysis were suggested to improve the power[7,8].

#### Expected LMM test statistic for Unrelated+GRM

The expected LMM test statistic for scenario Unrelated+GRM (Table 1) was previously derived analytically under assumption of polygenic inheritance and in the absence of population structure and other artifacts[9–11]. Here we outline the derivation and show the connection with our multiplier $\gamma_\beta$.

Consider a set of $N$ unrelated individuals, $M_e$ effective unlinked genetic variants and the heritability $\sigma_g^2$ explained by variants. The expected statistic from Equations (4) and (7), averaged over *all* variants, is

$$\mathbb{E}(s_{LR}) = 1 + N\mathbb{E}(\beta^2) = 1 + N\sigma_g^2/M_e \tag{53}$$

$$\mathbb{E}(s_{LMM}) = 1 + \gamma_G \mathbb{E}(\beta^2) = 1 + \gamma_G N\sigma_g^2/M_e \tag{54}$$

The polygenic effect of variants is introduced as a random effect in LMM such that $\Sigma_y = \sigma_g^2 G + (1-\sigma_g^2)I$, where $G = WW^T/M$ is the genetic relationship matrix (GRM). Conditioning on the polygenic effect in LMM association tests increases power over LR tests, as the gain factor $\gamma_G$ is always greater than one[10].

$$\begin{aligned} \gamma_G &= 1/(1-r^2\sigma_g^2) \\ r^2 &= (1+\theta) - \sqrt{(1+\theta)^2 - 4\sigma_g^2\theta}/(2\sigma_g^2) \text{ where } \theta = N\sigma_g^2/M_e \\ r^2 &\approx N\sigma_g^2/M_e \text{ when } M_e > N \end{aligned} \tag{55}$$

When $M$ variants are independent, we then have $M_e = M$. Otherwise, the effective number of variants, $M_e$ can be estimated from data[10].

##### Relationship between $\gamma_G$ and multiplier $\gamma_\beta$ for Unrelated+GRM

The multiplier $\gamma_\beta$ for Unrelated+GRM in Equation (29) can be further simplified using the random matrix theory applied to the matrix $G$[11], which is a Wishart matrix. Using the Marchenko-Pastur distribution of the eigenvalues $\nu(x)$ of $G$, the integral for the density eigenvalues can be analytically computed and is directly related to $\gamma_G$ in Equation (55)[10,11].

$$\gamma_\beta(\text{Unrelated+GRM}) \approx \frac{1}{N} \int \frac{\nu(x)}{\sigma_a^2 x + \sigma_r^2} dx = \gamma_G \tag{56}$$

#### Data simulations

In power analysis of testing marginal genetic effect, we simulate trait mean as $\mathbb{E}(y) = \beta_g x_g$ on allelic (unstandardized) scale: the allelic effect size $\beta_g = 0.05$ and genetic variant $x_g$ has entries 0, 1, and 2 with minor allele frequency $p = 0.3$. The effect size $\beta_g = 0.05$ is allelic and corresponds to $\beta_{allele}$ in Equation (57). The genetic variant explains $\approx 0.1\%$ of trait variance.

Similarly in power analysis of testing gene-environment interaction effect, we simulate trait mean as $\mathbb{E}(y) = \beta_g x_g + \beta_e x_e + \beta_{ge} x_g * x_e$ on unstandardized scale. Genetic variant $x_g$ has etries 0, 1, and 2 with minor allele frequency $p = 0.3$. Binary exposure $x_e$ has entries 0 and 1 with frequency $f = 0.6$. All effect sizes, main genetic $\beta_g$, main environmental $\beta_e$ and interaction $\beta_{ge}$ are equal to 0.1. The gene-environment interaction (standardized) explains $\approx 0.1\%$ of trait variance.

In simulations of unrelated individuals under the polygenic model[10], we set the number of individuals $N = 1,000$, the number of all genetic variants $M = 2,000$ and the number of causal variants either $M_c = 200$ (default) or $M_c = 50$. We generate $M$ bi-allelic genetic variants with minor allele frequency $p = 0.5$, standardize them and store in a $N \times M$ matrix, $W$. We then generate a vector of genetic effect sizes, $b$, of causal variants from the normal distribution $\mathcal{N}(0, (\sigma_g^2/M_c)I)$, where $\sigma_g^2 = 0.8$ is heritability. Finally, trait is simulated as $y = Wb + \epsilon$, where the residual noise comes from the normal distribution $\mathcal{N}(0, (1 - \sigma_g^2)I)$. In the next step of fitting LMM to simulated data, we first construct GRM using either all or top associated variants (LR test statistics), then estimate variance components, in particular $\sigma_g^2$, by REML[10] and finally compute LMM test statistic. We note that we were able to fully recover the true heritability ($\approx 0.8$, $M_c = 200$), although the sample size of simulated dataset is relatively small ($N = 1,000$).

##### Standardized and allelic effect sizes

We use the standardized effect sizes for marginal genetic and gene-environment interaction effects, $\beta$ and $\delta$, respectively. The relation to allelic effect sizes can be derived through minor allele frequency of genetic variant, $p$, and, for example, frequency of binary environmental exposure, $f$[12].

$$\beta = 2p(1-p)\beta_{allele} \tag{57}$$

$$\delta = 2p(1-p)f(1-f)\delta_{allele} \tag{58}$$

The variance explained by genetic variant and gene-environment interaction is readily expressed through standardized effect sizes, $\beta^2$ and $\delta^2$, respectively.

#### Simulation results for Unrelated+GRM

Before studying the relative power between Unrelated and Unrelated+GRM scenarios on simulated data, we sought to examine the impact of several LMM configurations that differ by variant selection for GRM.

As described in Methods, we simulated a trait under the polygenic model on $N = 1,000$ unrelated individuals, $M = 2,000$ (unlinked) genetic variants and $M_c = 200$ causal variants that explain 80% of trait variance, heritability $\sigma_g^2$. We first performed association study by LR, from which we ranked variants by their association statistic. We then examined three sets of $M_s = 200$ selected variants ($M_s = M_c$): random variants (Random), top LR top associated variants (Top) and causal variants (Causal). When fitting LMM to estimate the heritability (a single run of data simulation), we observed that the three models revealed different estimates, $\hat{\sigma}_g^2$: 9% for Random, 65% for Top and 80% for Causal. The accuracy of recovering the true heritability was driven by the sample size, $N$, the number of selected variants for GRM, $M_s$ and the number of causal variants captured in GRM (Supplementary Figure S11). For three LMM configurations on Figure 1 the number of causal variants included in GRM was equal to 22 for Random, 90 for Top and 200 for Causal.

We then examined how LMM configurations with different sets of selected variants influenced estimation of the effective size multiplier, $\gamma_\beta$. LMM association statistics and the multiplier were computed by plugging the estimated heritability, $\hat{\sigma}_g^2$, and trait covariance, $\hat{\Sigma}_y$, into Equations (3), (4) and (29). Figure S9a shows that the effective size multiplier $\gamma_\beta$, derived using the proposed analytical formulation, accurately approximated empirical ratios between LR and LMM squared standard errors. Importantly, the approximation worked equally well for all three LMM configurations with different estimates of heritability and, consequently, trait covariance matrix.

We next evaluated the performance of the empirical effective size multiplier[13], which we denoted as $\gamma_e$ and described in [Supplementary Material][Expected statistic]. Figure S9b shows that the accuracy of the empirical multiplier $\gamma_e$ was variable across LMM configurations and dependent of sets of variants used in ratios. Given that the choice of top statistic is subjective, we explored two approaches to select top associated statistic for $\gamma_e$ in each LMM configuration: significant variants ($P < 1 \times 10^{-5}$ in LMM) and top variants (significant in LMM, $P < 1 \times 10^{-5}$, and nominally significant in LR, $P < 0.05$).

For first LMM configuration with random variants in GRM (left panel on Figure S9b), the multiplier $\gamma_e$ is trivially equal to one (LMM $\approx$ LR), because most of the random variants were null and explained nearly zero heritability. For second LMM configuration with top associated variants in GRM (middle panel on Figure S9b), the empirical multiplier $\gamma_e$ is consistently lower than the effective sample size multiplier $\gamma_\beta$. The reason for this mismatch can be explained by the composition of top associated statistic for $\gamma_e$, where almost a half of variants are null with the ratio expected to be one. For the last LMM configuration with all causal variants in GRM (right panel of Figure S9b), the empirical multiplier $\gamma_e$ largely overestimated $\gamma_\beta$ for a set of top associated *causal* variants. Particular causal variants with low effect sizes (Supplementary Figure S12) were significant only in LMM, because the residual variance was remarkably reduced, as $\approx$ 80% heritability was explained by GRM. This overestimation was partially mitigated if nominally insignificant variants in LR ($P > 0.05$) are filtered out. If one uses median instead of mean to estimate the ratio of test statistic for $\gamma_e$, the the estimator is less prone to outliers that are nominally insignificant variants in LR ($P > 0.05$) (see Supplementary Figure S10).

### Supplementary Tables

**Table S1:** Relationship inference criteria based on estimating kinship coefficients ($\phi$) recommended by the authors of KING; see Table 1 in ref.[14]. To distinguish between parent–offspring and full-sibling pairs with the same expected kinship coefficient 1/4, any such pair with IBS0 $\leq$ 0.0012 is called parent-offspring, as performed in the original UK Biobank article[15].

| Relationship | $\phi$ | Inference criteria |
|---|---|---|
| Monozygotic twin | $\frac{1}{2}$ | $\phi > \frac{1}{2^3}$ |
| Parent–offspring | $\frac{1}{4}$ | $\phi \geq \frac{1}{2^{5/2}}$ & $\phi < \frac{1}{2^{3/2}}$ & IBS0 $\leq 0.0012$ |
| Full sibling | $\frac{1}{4}$ | $\phi \geq \frac{1}{2^{5/2}}$ & $\phi < \frac{1}{2^{3/2}}$ & IBS0 $> 0.0012$ |
| 2nd Degree | $\frac{1}{8}$ | $\phi \geq \frac{1}{2^{7/2}}$ & $\phi < \frac{1}{2^{5/2}}$ |
| 3rd Degree | $\frac{1}{16}$ | $\phi \geq \frac{1}{2^{9/2}}$ & $\phi < \frac{1}{2^{7/2}}$ |

**Table S2:** Relative power of association study in a UK Biobank subsample of related pairs (up to the 2nd degree).
*The ESS multiplier has a minimum value at a given value of heritability.

| Relationship | No. pairs | No. individuals | ESS multiplier, $\gamma_\beta$* | Variance explained (heritability), $\sigma_a^2$* |
|---|---|---|---|---|
| Monozygotic twin | 179 | 358 | 0.500 | 1.000 |
| Parent–offspring | 6,273 | 11,202 | 0.922 | 0.560 |
| Full sibling | 22,664 | 41,512 | 0.929 | 0.531 |
| 2nd Degree | 11,115 | 20,196 | 0.982 | 0.511 |
| All above (<2nd Degree) | 40,231 | 68,910 | 0.939 | 0.537 |

### Supplementary Figures

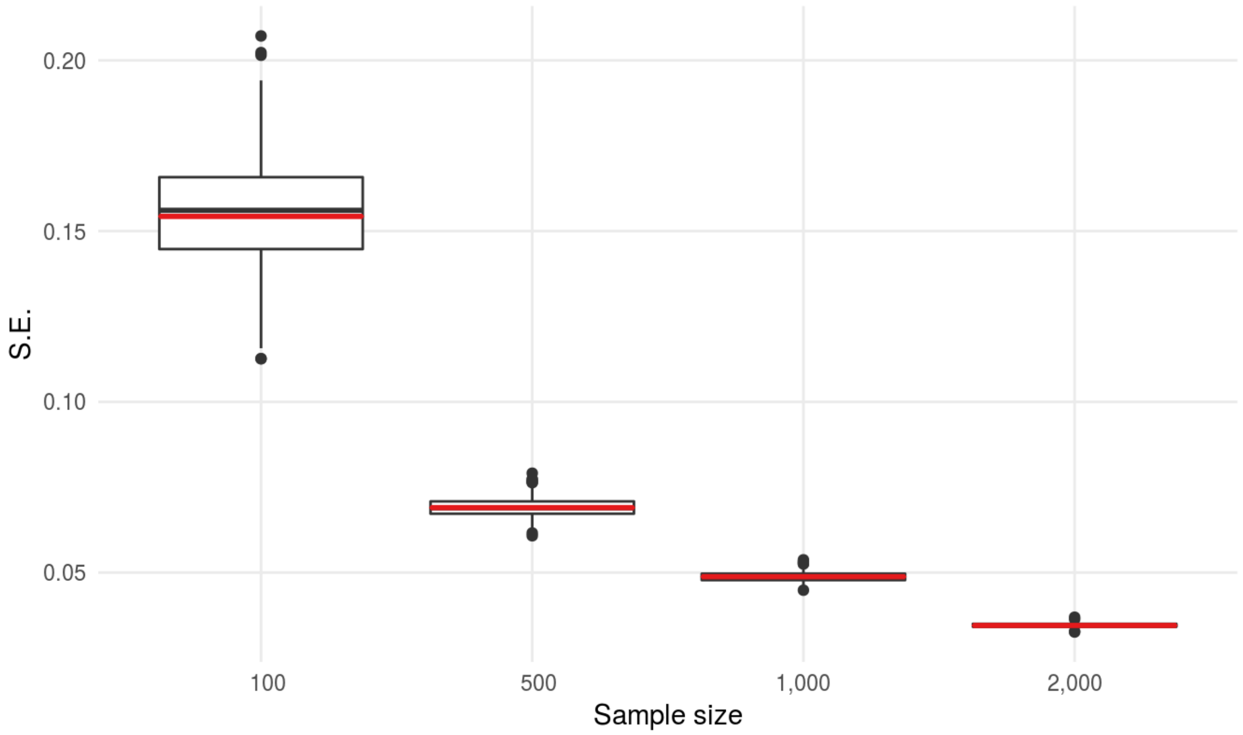


**Figure S1:** Simulation results in unrelated individuals when testing marginal genetic effect by LR. Association model to test the genetic effect $\beta$ is: $y \sim \mathcal{N}(w\beta, \Sigma_y = \sigma_r^2 I)$. Boxplots depict the distribution of empirical standard error of $\hat{\beta}$ (100 repetitions); the red line shows the analytical estimate of standard error of $\hat{\beta}$.

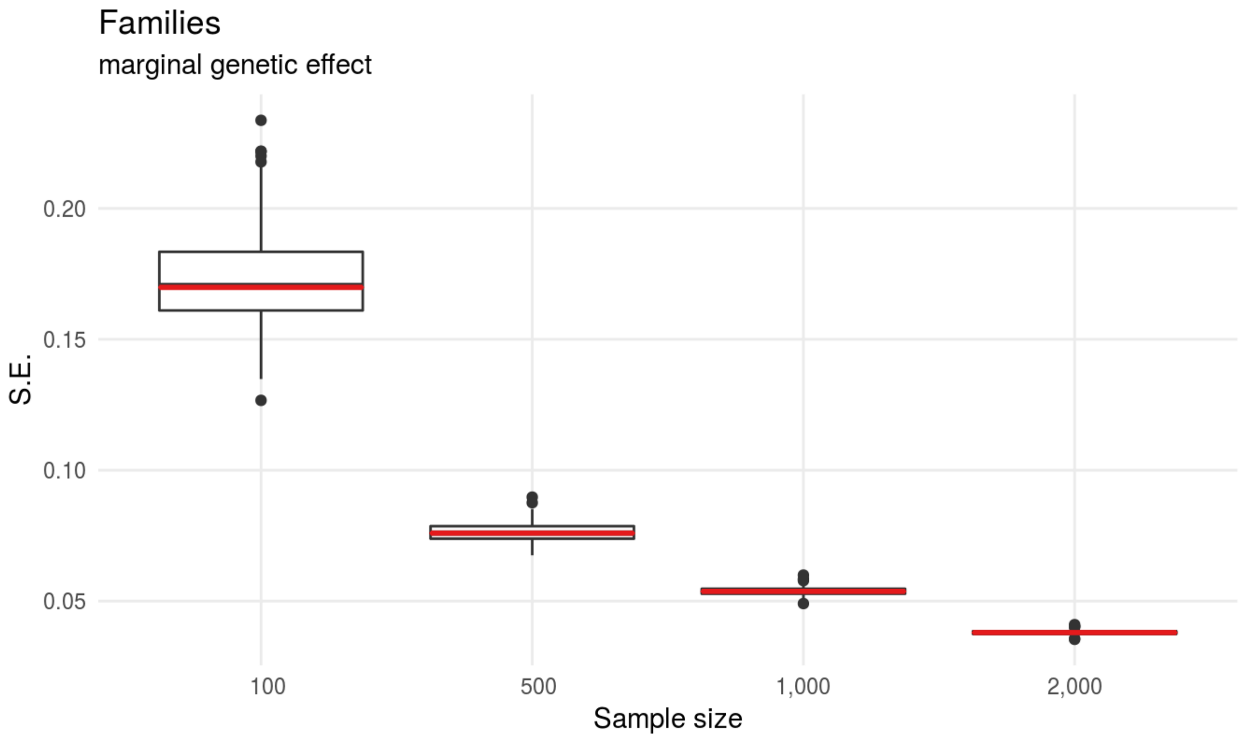


**Figure S2:** Simulation results in related individuals (nuclear families) when testing marginal genetic effect by LMM. Association model to test the genetic effect $\beta$ is: $y \sim \mathcal{N}(w\beta, \Sigma_y = \sigma_a^2 K + \sigma_r^2 I)$. Note that $cov(w) \equiv \Sigma_w = K$. Boxplots depict the distribution of empirical standard error of $\hat{\beta}$ (100 repetitions); the red line shows the analytical estimate of standard error of $\hat{\beta}$.

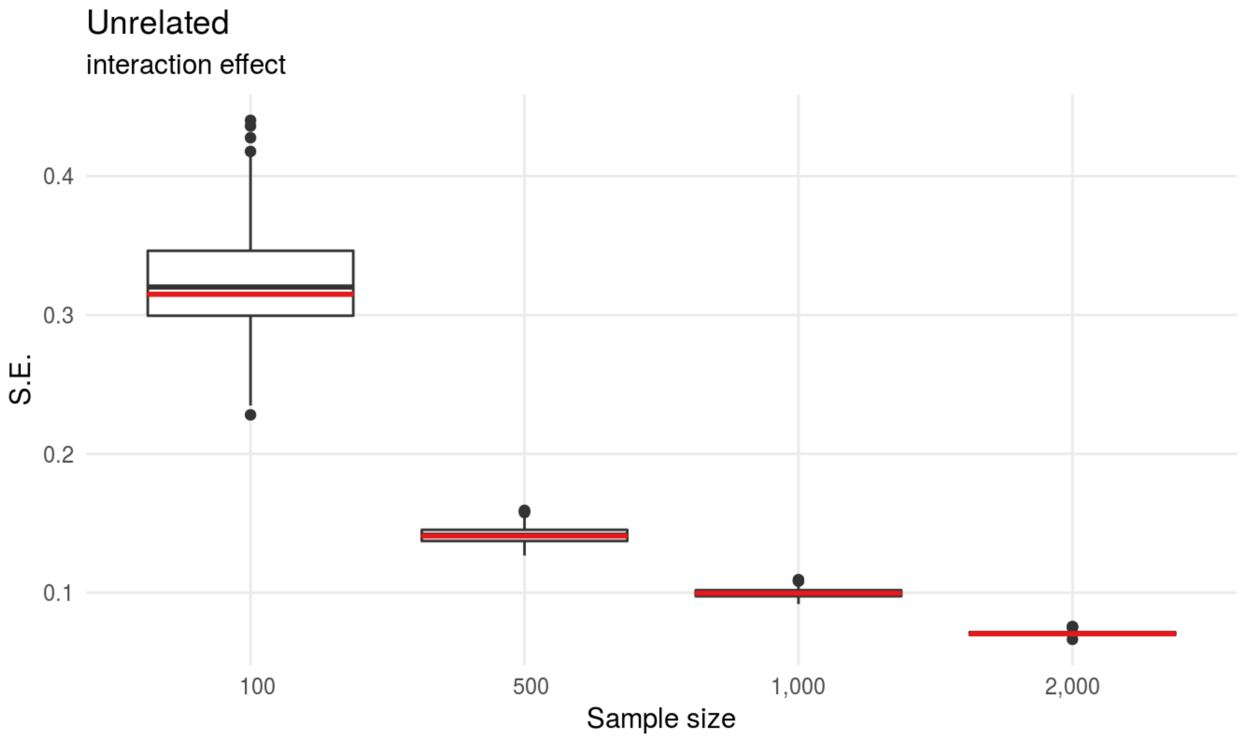


**Figure S3:** Simulation results in unrelated individuals when testing gene-environment interaction effect by LR. Association model to test the gene-environment interaction effect $\delta$ is: $y \sim \mathcal{N}(w\beta + d\tau + v\delta, \Sigma_y = \sigma_r^2 I)$. Boxplots depict the distribution of empirical standard error of $\hat{\beta}$ (100 repetitions); the red line shows the analytical estimate of standard error of $\hat{\beta}$.

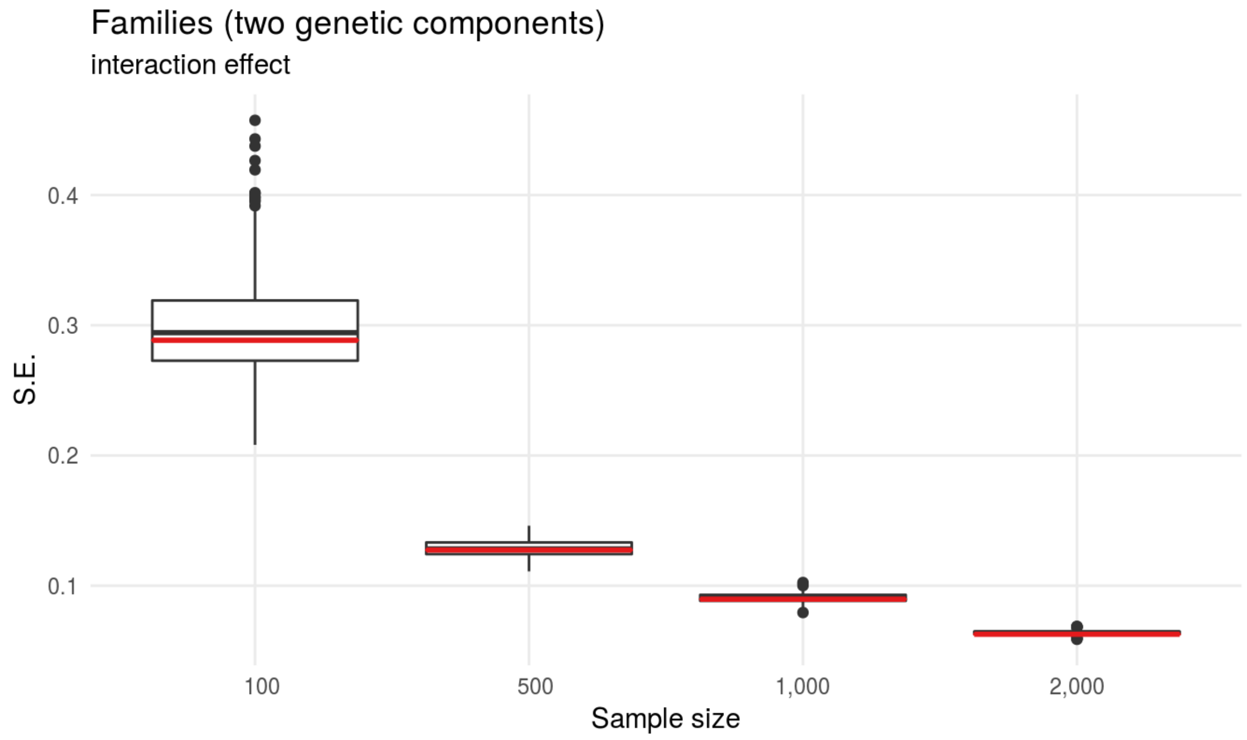


**Figure S4:** Simulation results in related individuals (nuclear families) when testing gene-environment interaction genetic effect by LMM. Association model to test the gene-environment interaction effect $\delta$ is: $y \sim \mathcal{N}(w\beta + d\tau + v\delta, \Sigma_y = \sigma_a^2 K + \sigma_{ai}^2 K_I + \sigma_r^2 I)$. Two genetic variance components, $\sigma_f^2$ and $\sigma_i^2$ are included, as described in[2]. Note that $cov(v) \equiv \Sigma_v = K_D$, Equation (20). Boxplots depict the distribution of empirical standard error of $\hat{\beta}$ (100 repetitions); the red line shows the analytical estimate of standard error of $\hat{\beta}$.

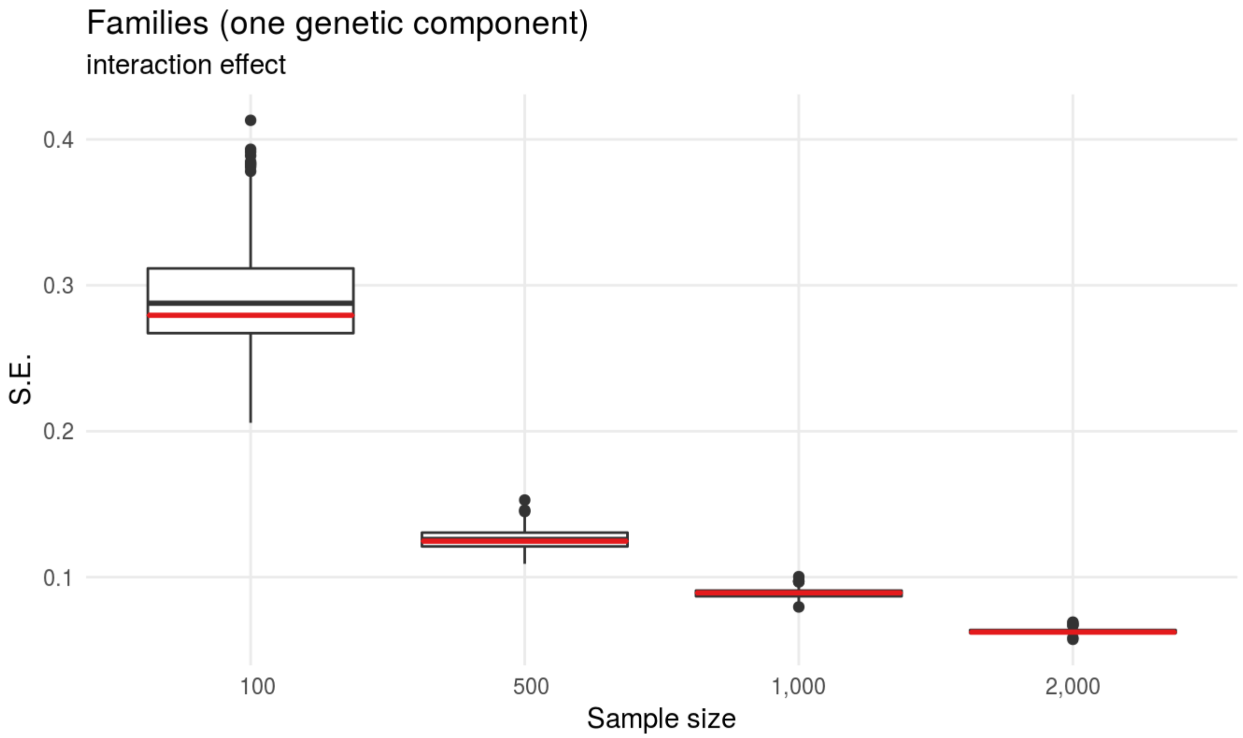


**Figure S5:** Simulation results in related individuals (nuclear families) when testing gene-environment interaction genetic effect by LMM. Association model to test the gene-environment interaction effect $\delta$ is: $y \sim \mathcal{N}(w\beta + d\tau + v\delta, \Sigma_y = \sigma_a^2 K + \sigma_r^2 I)$. Only one genetic variance component, $\sigma_f^2$, is included, in contrast to two genetic components described in[2]. Note that $cov(v) \equiv \Sigma_v = K_D$, Equation (20). Boxplots depict the distribution of empirical standard error of $\hat{\beta}$ (100 repetitions); the red line shows the analytical estimate of standard error of $\hat{\beta}$.

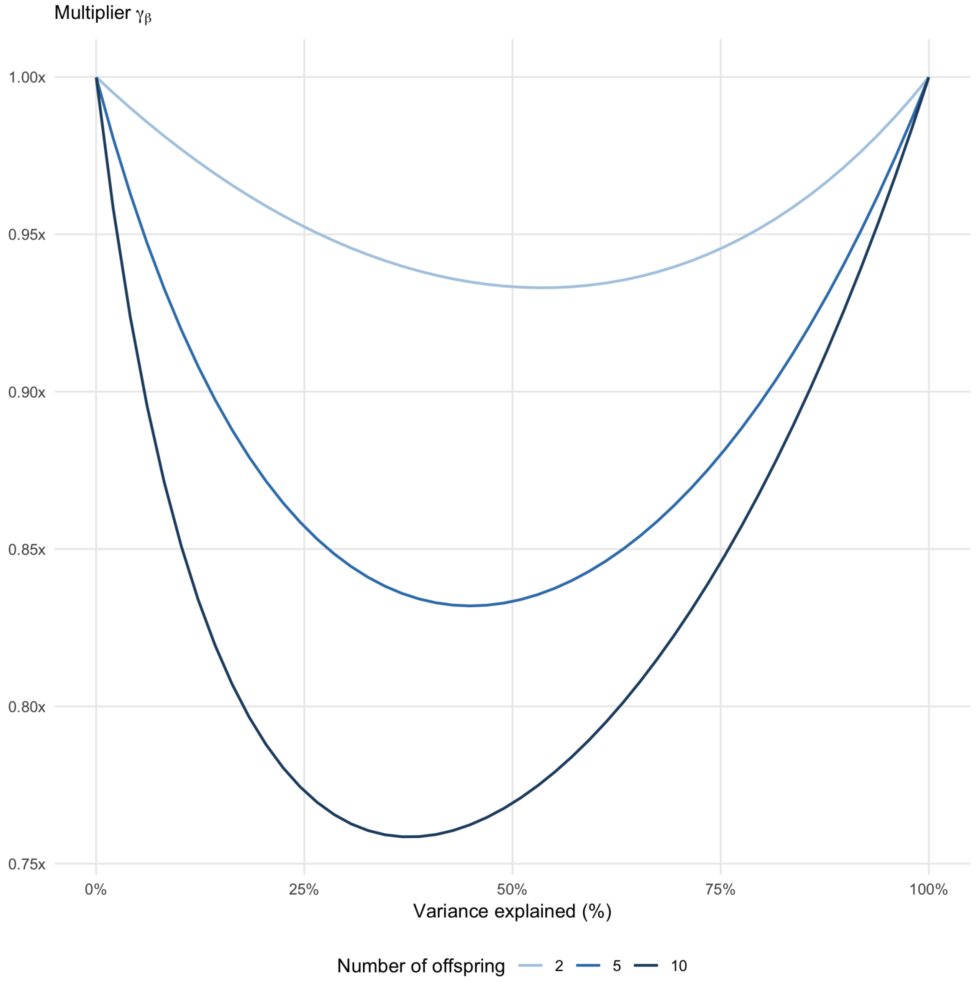


**Figure S6:** The effective size multiplier $\gamma_\beta$, analytically computed for nuclear families (2 parents and offspring), varies with proportion of variance explained by family relationships (heritability $\sigma_a^2$) and family structure (the number of offspring in nuclear families).

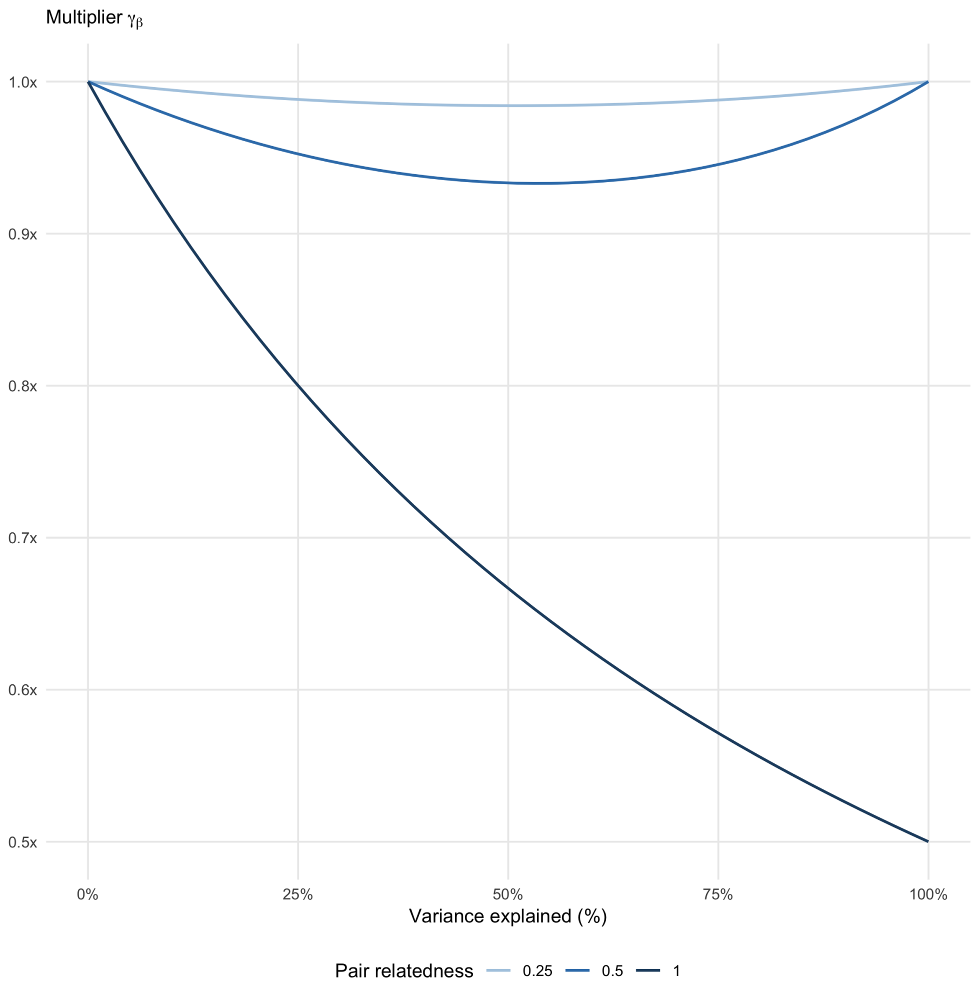


**Figure S7:** The effective size multiplier $\gamma_\beta$, analytically computed for related pairs, varies with proportion of variance explained by family relationships (heritability $\sigma_a^2$) and family structure (pair relatedness). The relatedness for different pairs (the double kinship coefficient): 0.125 for cousins, 0.5 for siblings, and 1 for monozygotic twins.

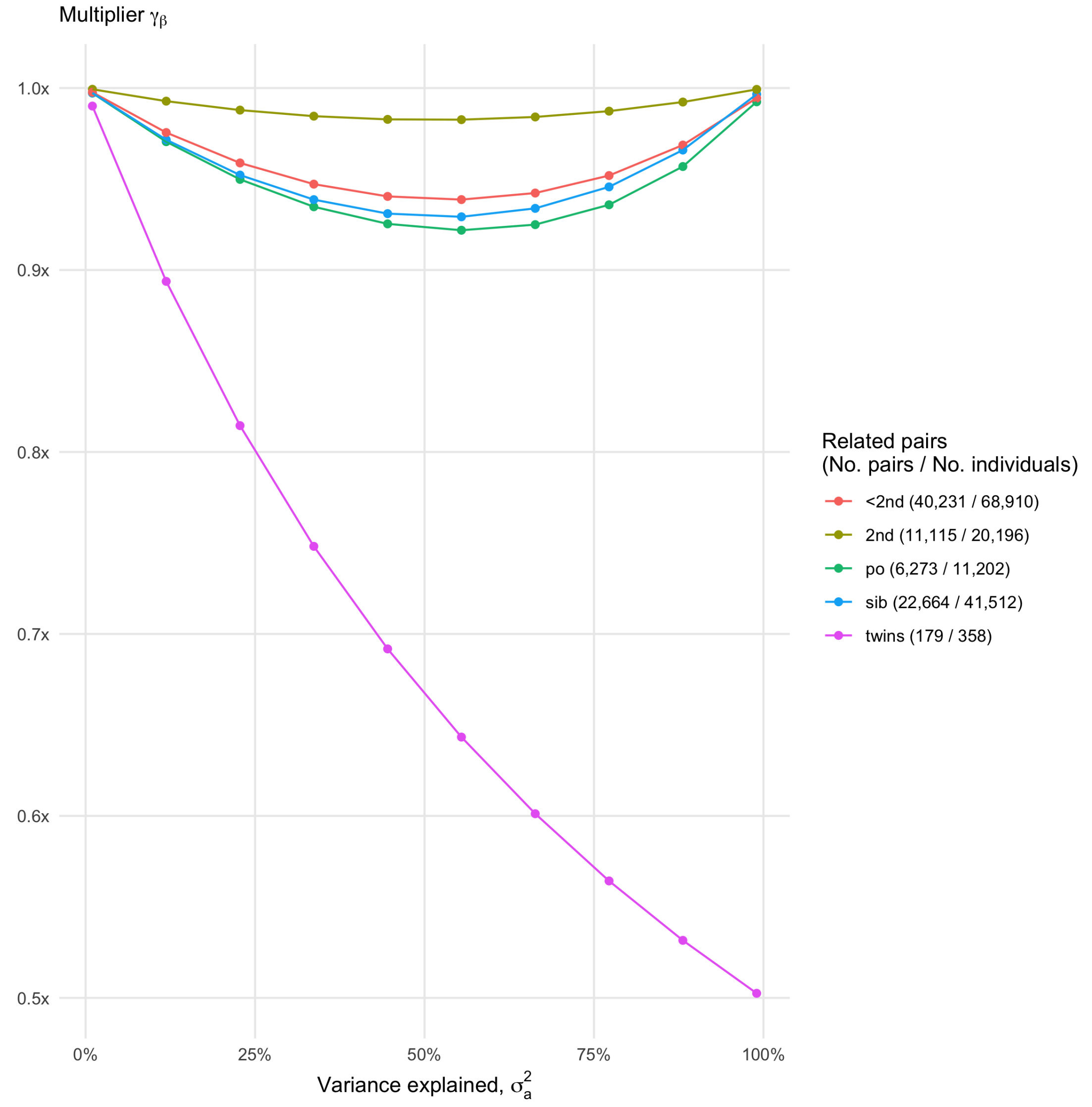


**Figure S8:** The effective size multiplier $\gamma_\beta$ is analytically estimated in 68,910 UK Biobank unrelated individuals (up to 2nd degree; see Supplementary Table S2). The multiplier is a function of the variance explained by family relationships (heritability $\sigma_a^2$) and the strength of genetic relatedness (twins, monozygotic twins: 1; sib, sibling pairs: 0.5; po, parent-offspring pairs: 0.5; 2nd, 2nd-order relatives: 0.125).

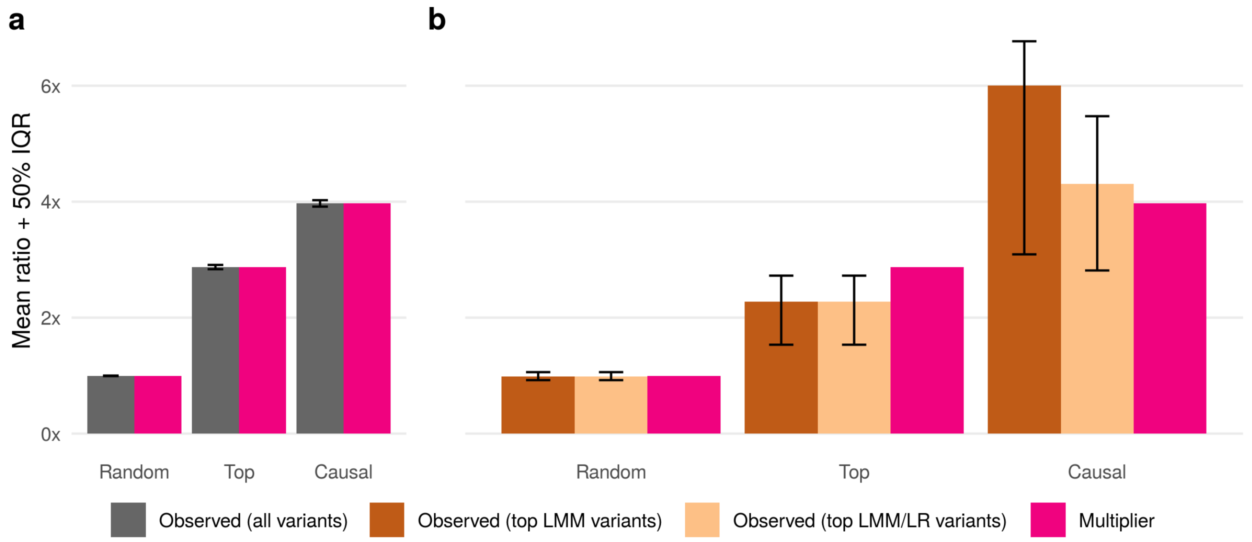


**Figure S9:** Validation of the analytical multiplier $\gamma_\beta$ on simulated data under Unrelated+GRM and Unrelated scenarios (N = 1,000, M = 2,000, $M_c$ = 200 and $M_s$ = 200; see Data simulations in Supplementary Material). Three sets of $M_s$ = 200 variants are selected to build GRM in LMM: random variants (Random), top LR top associated variants (Top) and causal variants (Causal). (a) The effective size multiplier $\gamma_\beta$ (red bars) accurately approximates empirical ratios of squared standard errors (dark grey bars) for every set of $M_s$ variants used in GRM. (b) The empirical multiplier $\gamma_e$ (brown and beige bars) is computed at different sets of variants: significant variants ($P_{LMM} < 1 \times 10^{-5}$ in LMM) and top variants (significant in LMM, $P_{LMM} < 1 \times 10^{-5}$, and nominally significant in LR, $P_{LR} < 0.05$). The multipliers $\gamma_e$ and $\gamma_\beta$ match only in the trivial case when random variants in GRM capture nearly zero heritability (Unrelated $\approx$ Unrelated+GRM). Otherwise, $\gamma_e$ gives biased estimates. Heights of dark grey, brown and beige bars represent mean values, while error bars range from 1st to 3rd quartiles. The multiplier $\gamma_e$ on panel (b) is not reported for sets of all variants (dark grey bars on panel (a)), because the mean statistic is not robust to outliers, which are causal variants with low effect sizes (significant in LMM and insignificant in LR). See also Supplementary Figure S10 for reported median ratios and Supplementary Figure S12 for distribution of tests statistics at causal variants, including causal variants with low effect sizes.

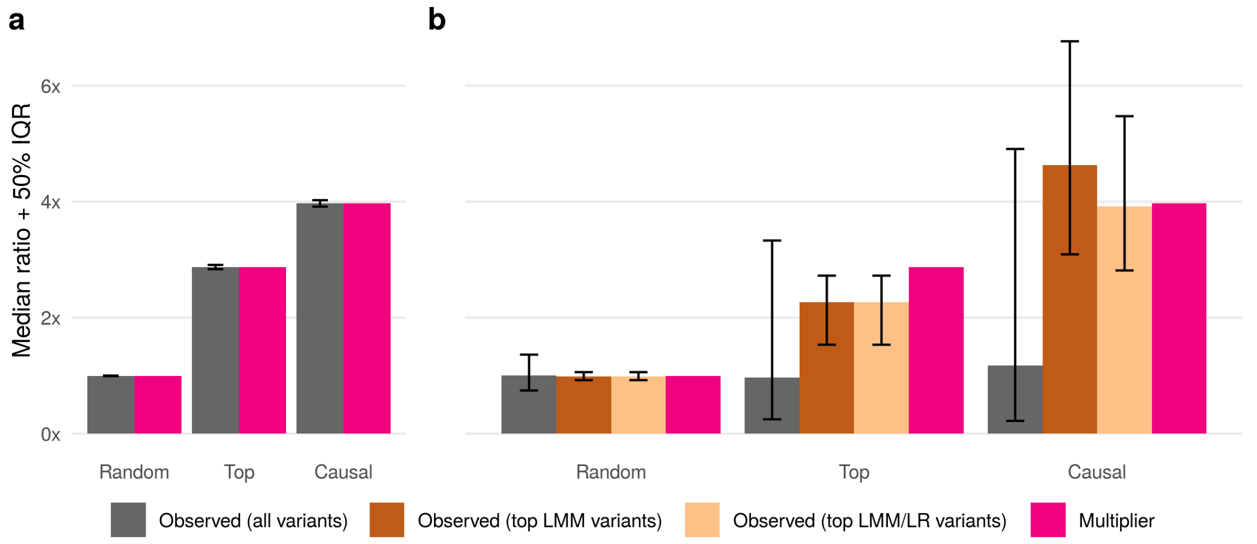


**Figure S10:** Results on simulated data, reported on Figure S9, using median rather than mean in computing (a) ratios of squared standard errors and (b) ratios of squared test statistic. Because of using median, the multiplier $\gamma_e$ on panel (b) for sets of all variants (dark grey bars) are reported.

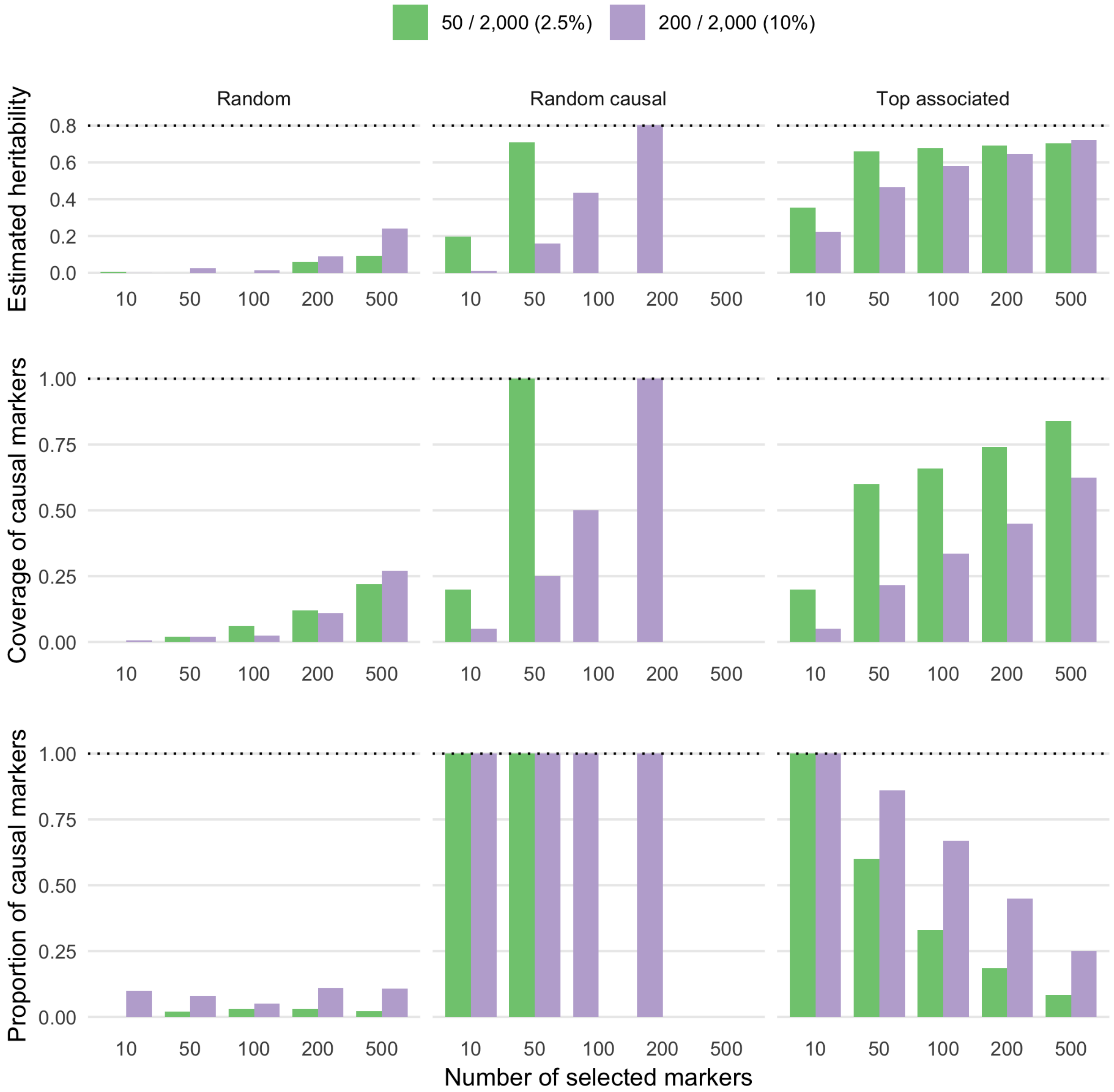


**Figure S11:** Estimated heritability is reported on simulated data under Unrelated+GRM scenario (N = 1,000, M = 2,000, $M_c$ = {50, 200} and $M_s$ = {10, 50, 100, 200, 500}; see Data simulations in Supplementary Material). Three sets of $M_s$ variants are selected to build GRM in LMM: random variants (Random), random causal variants (Causal) and top LR top associated variants (Top).

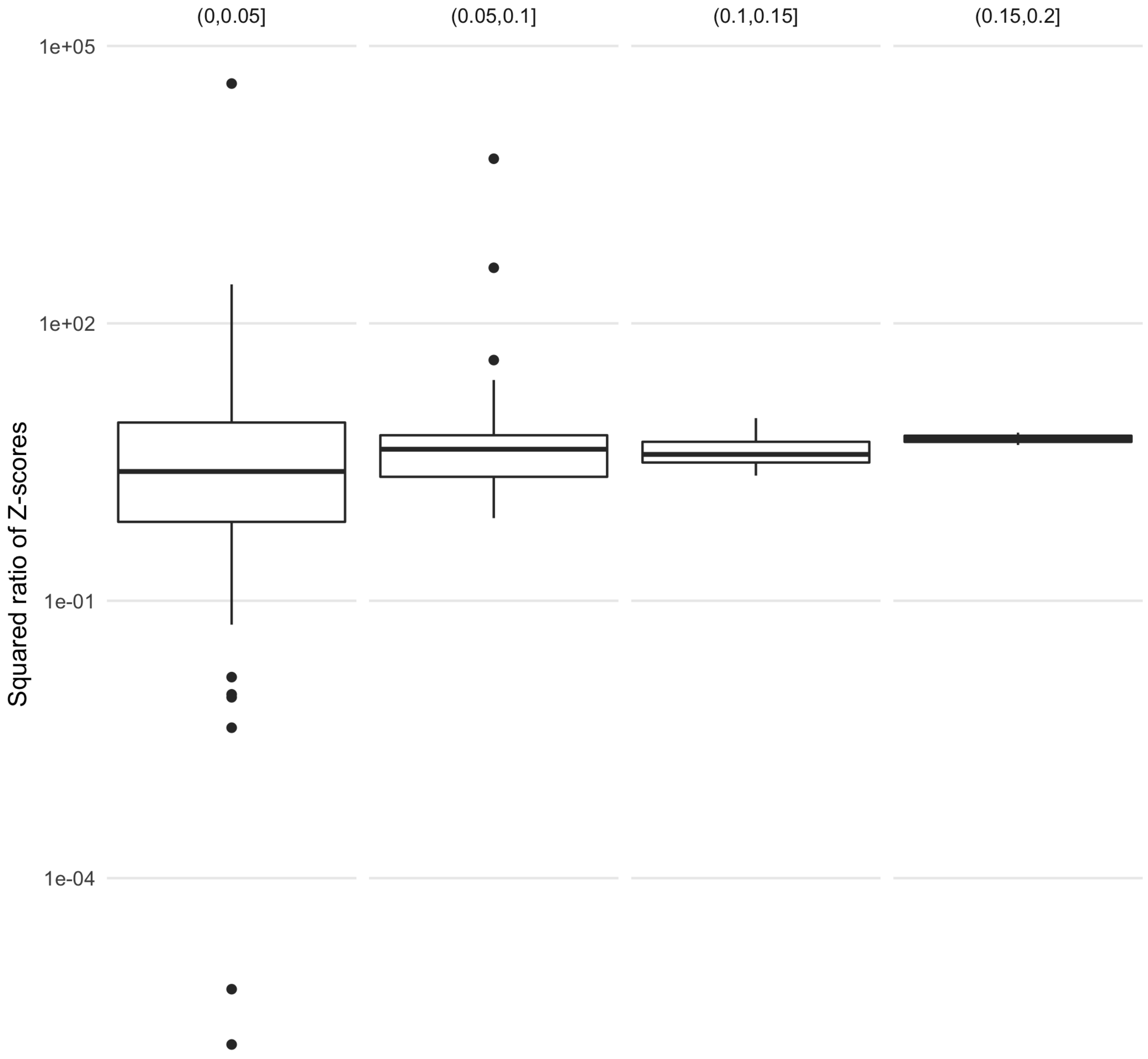


**Figure S12:** Distribution of LMM and LR test statistics (Z-scores) on simulated data (N = 1,000, M = 2,000, $M_c$ = 200 and $M_s$ = 200; see Data simulations in Supplementary Material). Ratios of association statistics LMM/LR between Unrelated and Unrelated+GRM scenarios are computed at causal variants and stratified by the effect sizes. Outlier points above the 75% quantile of box plots correspond to causal variants with low effect sizes that are insignificant in LR, but become significant in LMM. These particular variants inflates the empirical multiplier $\gamma_e$ computed on a set of causal variants (Figure S9). All causal variants are included in GRM when producing LMM test statistics.

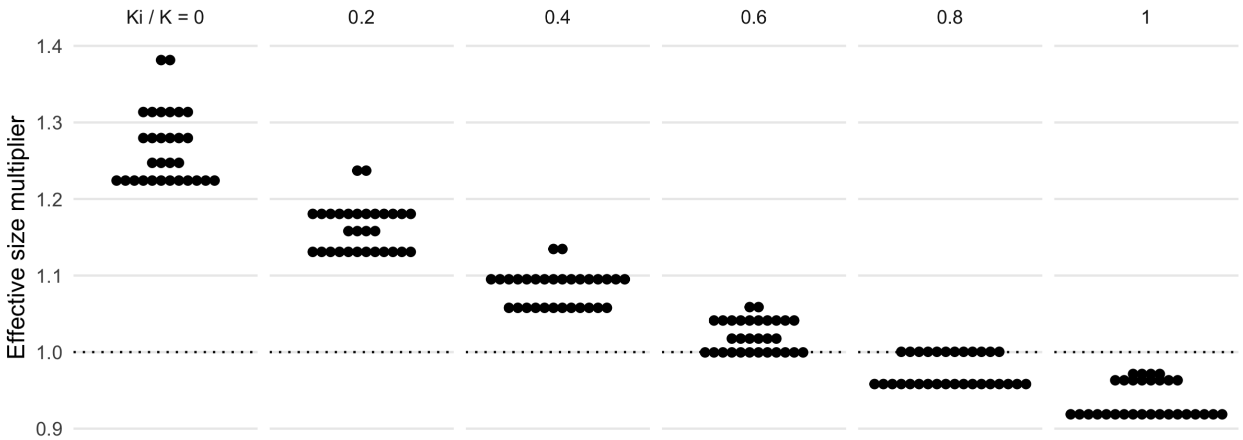


**Figure S13:** The effective size multiplier for gene-environment interaction effect $\gamma_\delta$ is analytically computed for nuclear families with 2 parents and 3 offspring. All possible realizations of a binary exposure within a family are considered, that results in different exposure frequencies. The distribution of $\gamma_\delta$ is shown as a dotplot for six combinations of variance components in LMM. Recall that the association model to test the gene-environment interaction effect $\delta$ is: $y \sim \mathcal{N}(w\beta + d\tau + v\delta, \Sigma_y = \sigma_a^2 K + \sigma_{ai}^2 K_I + \sigma_r^2 I)$. Each panel has its own ratio $\sigma_{ai}^2 / \sigma_a^2$, for instance, $\sigma_{ai}^2 = 0$ on the left panel and $\sigma_{ai}^2 = \sigma_a^2$ on the right panel.

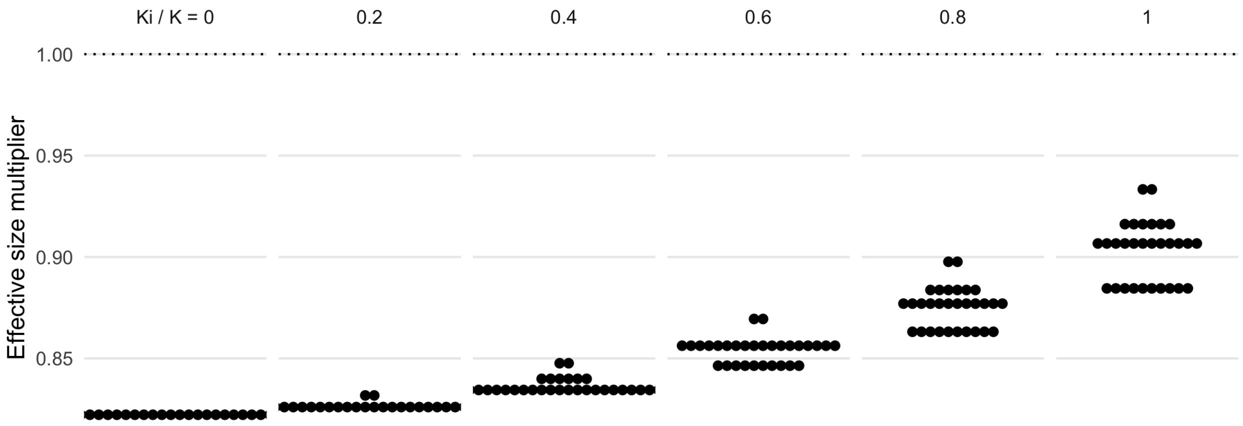


**Figure S14:** The effective size multiplier for genetic effect $\gamma_\beta$ (instead of $\gamma_\delta$ for gene-environment interaction effect) is analytically estimated on the same nuclear family data as on Figure S13.
